# Monoclonal antibodies targeting the FimH adhesin protect against uropathogenic *E. coli* UTI

**DOI:** 10.1101/2024.12.10.627638

**Authors:** Edward D. B. Lopatto, Jesús M. Santiago-Borges, Denise A. Sanick, Sameer Kumar Malladi, Philippe N. Azimzadeh, Morgan W. Timm, Isabella F. Fox, Aaron J. Schmitz, Jackson S. Turner, Shaza Sayed Ahmed, Lillian Ortinau, Nathaniel C. Gualberto, Jerome S. Pinkner, Karen W. Dodson, Ali H. Ellebedy, Andrew L. Kau, Scott J. Hultgren

**Affiliations:** Department of Molecular Microbiology, Washington University in St Louis, St Louis, MO, U.S.A; Center for Women’s Infectious Disease Research, Washington University in St Louis, St Louis, MO, U.S.A; Division of Allergy and Immunology, Department of Medicine, Washington University in St Louis, St Louis, MO, U.S.A; Department of Pathology and Immunology, Washington University School of Medicine, St Louis, MO, U.S.A; Center for Vaccines and Immunity to Microbial Pathogens, Washington University School of Medicine, St. Louis, MO, U.S.A

## Abstract

As antimicrobial resistance increases, urinary tract infections (UTIs) are expected to pose an increased burden in morbidity and expense on the healthcare system, increasing the need for alternative antibiotic-sparing treatments. Most UTIs are caused by uropathogenic *Escherichia coli* (UPEC), while *Klebsiella pneumoniae* causes a significant portion of non-UPEC UTIs. Both bacteria express type 1 pili tipped with the mannose-binding FimH adhesin critical for UTI pathogenesis. We generated and biochemically characterized 33 murine monoclonal antibodies (mAbs) to FimH. Two mAbs protected mice from *E. coli* UTI. Mechanistically, we show that this protection is Fc-independent and mediated by the ability of these mAbs to sterically block FimH function. Our data reveals that FimH mAbs hold promise as an antibiotic-sparing treatment strategy.

## Introduction

Urinary tract infections (UTIs) affect over 400 million individuals worldwide yearly (*1*, *2*), leading to $2.8 billion in healthcare and productivity-related costs annually in the US alone (*3*). Around 25% of individuals will suffer from recurrent UTIs which severely impairs their quality of life (*4*). Further, 27% of all sepsis cases can be traced to urinary origin (*5*). This is aggravated by the increased prevalence of multidrug-resistant uropathogens (*6*), such as uropathogenic *Escherichia coli* (UPEC), responsible for 70-90% of UTIs, and *Klebsiella pneumoniae*, one of the most prevalent non-UPEC uropathogens (*7–9*). UTIs represent the fourth leading cause of death attributed to or associated with antibiotic resistance (*10*). Thus, developing antibiotic-sparing strategies to prevent UTIs caused by these difficult-to-treat uropathogens is crucial. One promising approach for new antibacterial treatments is to neutralize key extracellular adhesins to prevent bacterial colonization and invasion into tissue and biofilm formation.

UPEC and *K. pneumoniae* express chaperone-usher pathway (CUP) type 1 pili that are tipped with the mannose-binding FimH adhesin essential in **i)** bladder colonization, **ii)** ascension to cause pyelonephritis, **iii)** invasion into terminally differentiated umbrella cells of the bladder, **iv)** the formation of intracellular biofilms in luminal bladder cells, and **v)** causing an epigenetic imprint in the bladder that predisposes to recurrent UTI (*11–18*). FimH is a two-domain protein with an N-terminal lectin domain containing a deep binding pocket that recognizes mannose with stereochemical specificity (*19*) and a C-terminal pilin domain linking the adhesin to the pilus. At the tip of the assembled type 1 pilus, FimH exists in a conformational equilibrium between a high-affinity relaxed state and a low-affinity tense state controlled by structural interactions between the FimH lectin and pilin domains (*20*). In the high-affinity relaxed state, the FimH lectin domain is highly mobile with respect to the pilin domain. In contrast, in the low-affinity tense state, the pilin domain constrains the lectin domain and allosterically deforms the mannose-binding pocket. UPEC FimH occupies both tense and relaxed conformations whereas the equilibrium in the highly invariant and conserved *K. pneumoniae* FimH is primarily shifted towards the tense low-binding conformation, explaining its poor mannose binding properties despite an identical mannose binding site (*21*).

A vaccine against UPEC FimCH has revealed an 73% reduction in recurrent UTIs caused by UPEC or *Klebsiella* spp. in a phase 1A/1B clinical trial, showing potential to prevent the two most common UTI pathogens with a FimH targeted therapeutic (*22*, *23*). The effectiveness of FimH vaccination is associated with antibody responses that inhibit FimH binding (*24*, *25*). Here, we characterized monoclonal antibodies (mAb) from mice immunized with *E. coli* and *K. pneumoniae* FimH lectin domains and discovered cross-reactive antibodies that bind to four distinct FimH structural epitopes (**Class 1-4)**, which block FimH binding *in vitro*. Using cryo-EM, we discovered that the mAbs are selectively bound to the epitopes displayed in the high-affinity relaxed conformation of FimH. Using binding studies and mouse UTI models, we identified **Class 1** mAbs that blocked FimH binding to mannose through steric interference leading to protection against UPEC in mouse UTI models. From structure to an antibiotic alternative therapeutic, these results guide future optimization of FimH mAbs and vaccination strategies to treat *E. coli* and *K. pneumoniae* UTIs, two of the most prominent uropathogens with increasing antibiotic resistance.

## Results

### FimH_LD_ mAbs can cross-react with E. coli and K. pneumoniae FimH

Female C57BL/6J mice were immunized with the lectin domain truncates of FimH (FimH_LD_) from either *E. coli* (UTI89) or *K. pneumoniae* (TOP52). Heavy chain V-D-J and light chain V-J fragments from sorted plasmablasts isolated from draining lymph nodes were cloned into human IgG expression vectors to create chimeric murine/human mAbs as previously described (*26*). In total, 33 clonally distinct mAbs were generated from *E. coli* (8 mAbs) and *K. pneumoniae* (25 mAbs) that bound the respective FimH_LD_ (referred to as Ec or Kp mAbs respectively). The FimH_LD_ structures of *E. coli* and *K. pneumoniae* are highly homologous (RMSD = 0.420) with an 86% amino acid sequence identity, including an identical binding pocket (Fig. 1A-B, S1). By ELISA, most mAbs bound to their respective FimH_LD_ antigen with half-maximal effective binding concentrations (EC_50_) ranging from 8 to 50 ng/mL, while some bound more weakly with EC_50_ values from 50 to 751 ng/mL (Figure 1D). Half (4 of 8) of the Ec mAbs also reacted with *K. pneumoniae* FimH_LD_ and 80% (20 of 25) of Kp mAbs reacted with *E. coli* FimH_LD_. Further, eight mAbs (2 Ec and 6 Kp mAbs) that reacted with both FimH antigens also bound with high-affinity to a third structurally similar (RMSD = 0.699 to Ec FimH) *E. coli* adhesin, FmlH lectin domain (FmlH_LD_), an adhesin that binds to exposed galactose residues on bladder tissue during chronic cystitis infections (*27*) (Fig. 1C-D, S1).

**Fig. 1.**
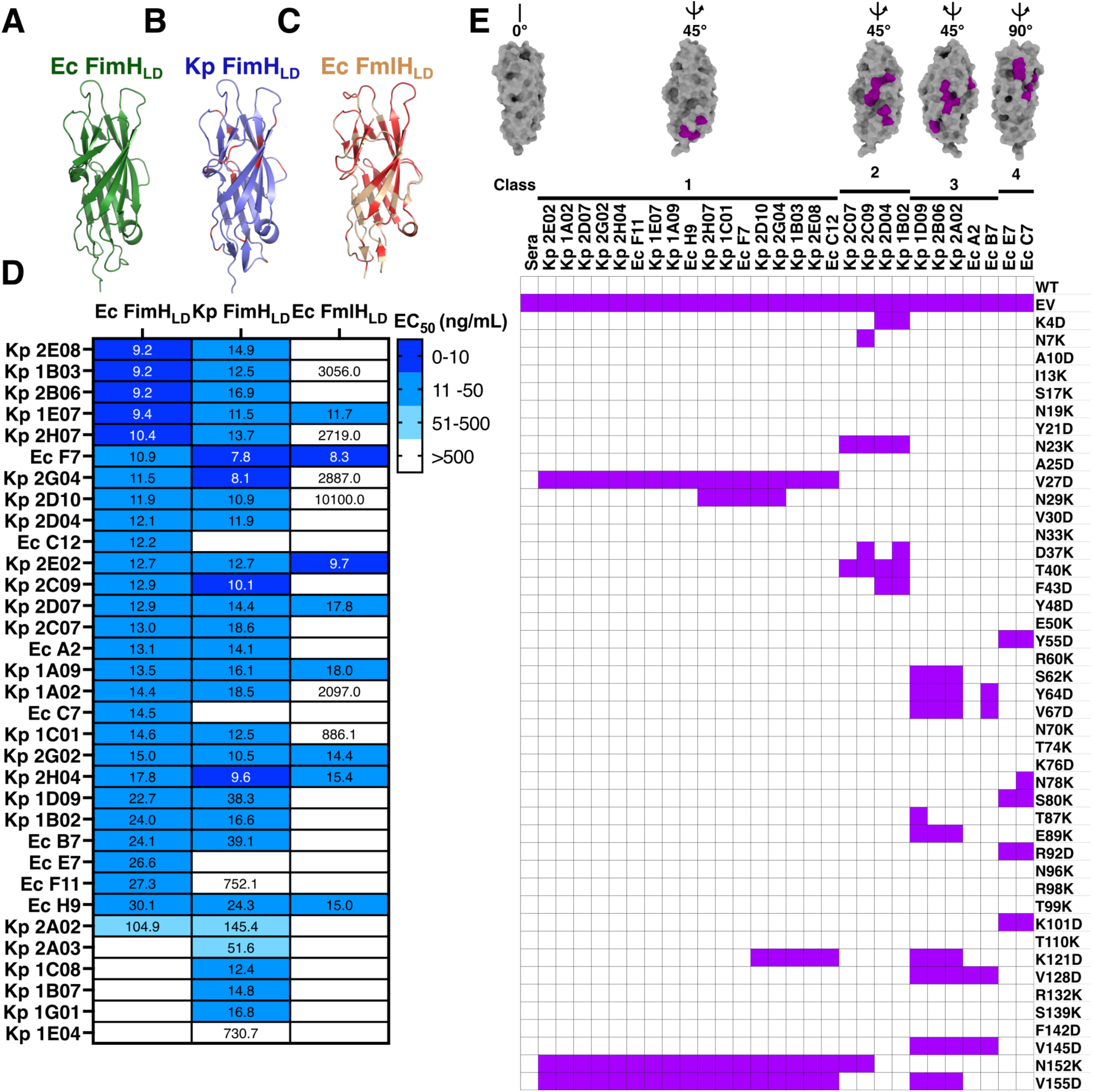
FimH mAbs bind to four distinct epitope classes. Structures of (**A**) *E. coli* FimH_LD_ (PDB 1KLF), (**B**) *K. pneumoniae* FimH_LD_ (PDB 9AT9), and (**C**) *E. coli* FmlH_LD_ (PDB 6AOW). Residue differences from *E. coli* FimH_LD_ are highlighted in red. (**D**) ELISA EC_50_ values for each mAb to the listed protein. White cells with no values indicate EC_50_ values were above the range measured in the assay. (**E**) Epitope mapping of mAbs (top labels) to a panel of FimH mutants (right labels). Binding classes were determined by shared residues that abrogated mAb binding which are highlighted in purple on the surface of *E. coli* FimH_LD_ (above) (PDB 1KLF) and the table (below).

### FimH_LD_ mAbs bind diverse epitopes

To determine the structural regions of FimH_LD_ that were recognized by each mAb, we generated a mutant library consisting of 45 surface-exposed mutations to bulky charged residues across the lectin domain of full-length *E. coli* FimH. Binding of the mAbs to the stable mutant FimH proteins was determined by ELISA, which revealed four binding site classes (Fig. 1E, structural regions of FimH are defined in Fig. S1 and Table S1). **Class 1** mAbs (4 Ec and 13 Kp mAbs), bind with high affinity to both *E. coli* and *K. pneumoniae* FimH_LD_ and FmlH_LD_. **Class 1** mAbs were unable to bind *E. coli* FimH **Class 1** epitope residue mutants V27D, N152K, and V155D located in the insertion and swing loops, and in the part of the linker between the pilin and receptor binding domains, suggesting they bind to the base of the lectin domain. **Class 2** mAbs (4 Kp mAbs) bound to *E. coli* and *K. pneumoniae* FimH_LD_, but did not react with FmlH_LD_, consistent with the sequence differences between *E. coli* FimH_LD_ and *E. coli* FmlH_LD_ (Fig. S1). **Class 2** mAbs shared the inability to bind to the **Class 2** epitope residue mutants N23K and T40K in *E. coli* FimH suggesting that they bind to the side of the FimH_LD_ body between the base of the binding clamp loop and basal swing loop. A majority of **Class 3** mAbs (2 Ec mAbs and 3 Kp mAbs), bound to both *E. coli* and *K. pneumoniae* FimH_LD_, but not to FmlH_LD_. **Class 3** mAbs were unable to bind to **Class 3** epitope residue mutants S62K, Y64D, V67D, E89K, K121D, V128D, V145D, and V155D in *E. coli* FimH, suggesting that they bind below the binding pocket to the opposite lateral side of the FimH_LD_ body from the **Class 2** epitope, covering regions on β-sheet B below the clamp loop and peripheral alpha-helix. **Class 4** mAbs (2 Ec mAbs) were only able to bind *E. coli* FimH_LD,_ and not to *K. pneumoniae* FimH_LD_ or *E. coli* FmlH_LD_ and were unable to bind to **Class 4** epitope residue mutants Y55D, S80K, R92D, and K101D in *E. coli* FimH, suggesting that they bound at the base of binding loop two and backside of FimH_LD_. Five Kp mAbs were not mapped as they did not bind *E. coli* FimH. Thus, immunization of *E. coli* and *K. pneumoniae* FimH_LD_ antigens generated mAbs bound to four unique surfaces (**Class 1-4** epitopes) on *E. coli* FimH. Below, mAb nomenclature denotes the epitope class recognized by the mAb after the Ec or Kp designation. Kp FimH_LD_ mAbs whose epitopes could not be determined are left unnumbered.

### Structural basis of FimH mAb recognition

The structural basis of **Class 1-3** mAbs binding to FimH_LD_ was determined by cryo-electron microscopy (cryo-EM) of FimCH complexed with fragment antigen-binding regions (Fabs) of Kp1 2H04, Ec1 F7, Kp2 2C07, and Ec3 B7 (Fig. 2 and S2-4). In each case, the observed interaction between FimH_LD_ and Fabs confirmed and extended the epitope mapping described above. The structural basis of Kp1 2H04 Fab and Ec1 F7 Fab binding to **Class 1** epitope of FimH revealed that they both interacted with the FimH_LD_ swing loop and linker regions (**Class 1** epitope residues A24 to N29 and N151 to T158) and coordinated multiple aromatic residues around FimH_LD_ **Class 1** residue P26 (Fig. 2A-C, F, and G). However, Kp1 2H04 and Ec1 F7 Fabs bound to the *E. coli* FimH_LD_ at differing angles. Ec1 F7 Fab was rotated ∼20 degrees relative to Kp1 2H04 Fab bound to FimH_LD_ (Fig. 2C and S4). The structure of the **Class 2** Kp2 2C07 Fab - FimH_LD_ complex revealed strong binding to the **Class 2** epitope residues Y21 to A27 and N151 to D153 (Fig. 2D and H). **Class 3** Ec3 B7 Fab bound to β-sheet B of *E. coli* FimH including **Class 3** residue Y64 (Fig. 2I and F), which has been identified as a “toggle switch” between tense and relaxed FimH conformation, with the residue undergoing a major solvent-accessible surface area change in the conformational transition (*28*). This binding mode suggests a role of our **Class 3** mAbs specifically binding and stabilizing the relaxed FimH conformation, as mutagenesis of **Class 3** residue Y64 abolished **Class 3** mAb binding in the epitope mapping (Fig. 1). Altogether, the high-resolution cryo-EM structures identified critical FimH interactions of the mAb epitope classes.

**Fig. 2.**
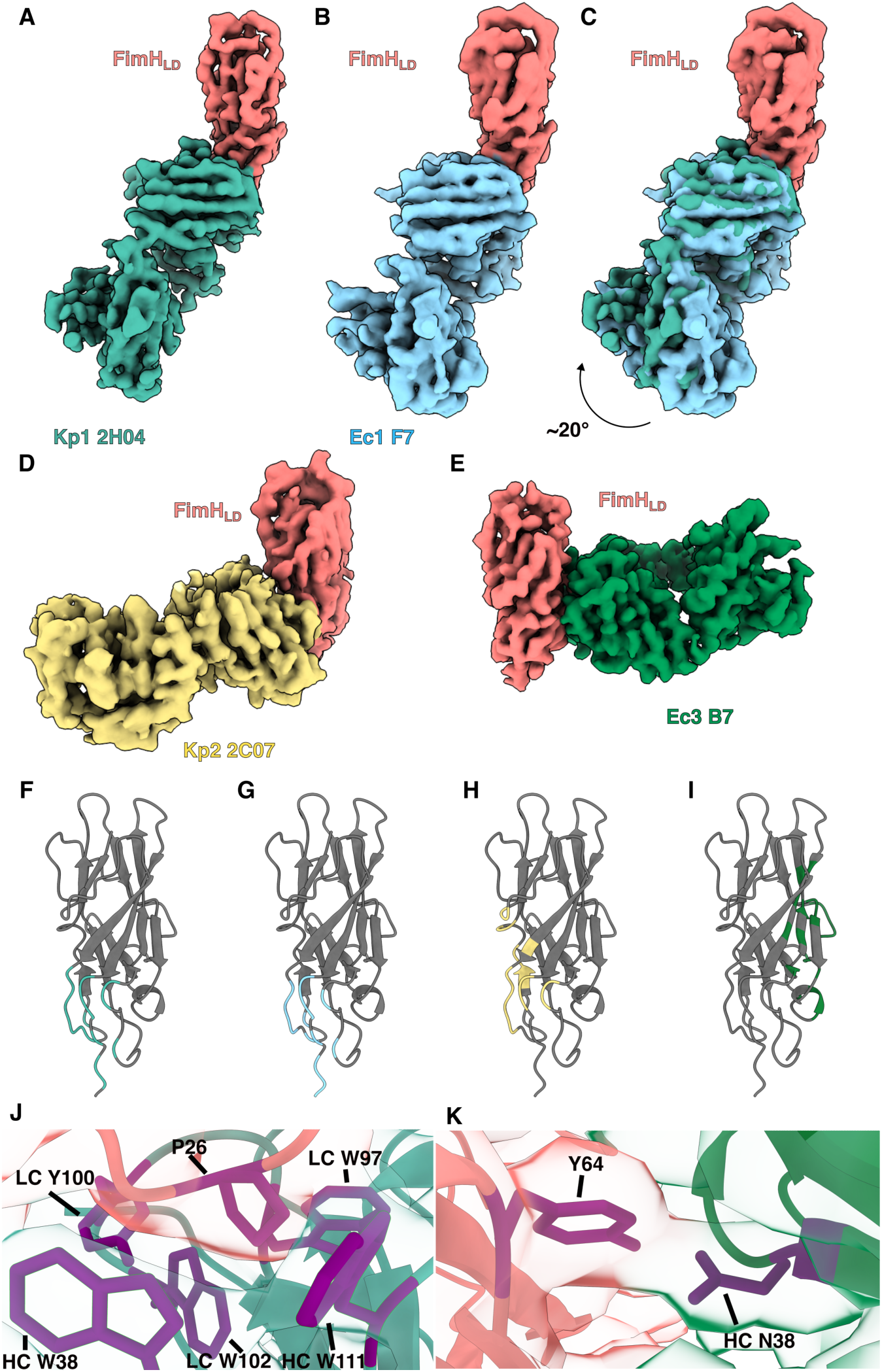
Structural basis of FimH mAb protection. (**A-E**) Cryo-EM density maps of Fabs bound to FimH_LD_. (**A**) Kp1 2H04 (teal), (**B**) Ec1 F7 (cyan), (**C**) Kp1 2H04 and Ec1 F7 maps superimposed on each other, (**D**) Kp2 2C07 (sand), and (**E**) Ec3 B7 (dark green) bound to FimH_LD_ (salmon). (**F-I**) The binding epitopes of the Fabs on FimH_LD_ colored by the same color scheme as panels A-E. Density map overlaid on model residue interactions of (**J**) Kp1 2H04 (cyan) with FimH P26 and (**K**) Ec3 B7 (dark green) with FimH Y64. mAb heavy chain residues are labeled “HC” and light chain residues are labeled “LC”.

### FimH_LD_ mAbs bind preferentially to the relaxed conformation

When FimH is incorporated at the tip of type 1 pili, the receptor binding domain samples a conformational equilibrium between low-affinity tense and high-affinity relaxed conformations (Fig. S1). The identified mAb binding epitopes of FimH_LD_, particularly **Class 1-3**, are in regions that vary extensively between tense and relaxed conformational states (*29*)(Fig S1, Table S1).

Thus, we investigated whether the conformational dynamics influence the epitopes recognized by **Class 1-4** mAbs by measuring binding to FimH-tipped piliated *E. coli* bacteria using ELISA. **Class 1** mAbs displayed the highest reactivity, while **Class 2-4** mAbs displayed greatly diminished reactivity to FimH tipped type 1 pili of *E. coli* (Fig. 3A). Thus, we tested **Class 1-4** mAbs against *E. coli* expressing conformational FimH variants. A majority of mAbs from **Classes 1-4** had increased binding to the relax-shifted A27V/V163A *E. coli* FimH mutant tipping type 1 pili and very weak binding to the tense-shifted A62S *E. coli* FimH mutant, despite similar levels of type 1 pili expression as measured by western blot (Fig. 3B, Fig. S5). To investigate binding of the highest affinity **Class 1-2** mAbs to FimH in a tip-like state, we used Bio-Layer Interferometry (BLI) to measure binding to recombinant FimH. We prepared “tip-like” recombinant full-length FimH by incubating *E. coli* FimCH with the N-terminal extension (Nte) peptide of FimG resulting in a donor strand exchange reaction where the FimG Nte displaces the FimC chaperone to produce *E. coli* FimG_nte_H complex that samples dynamic conformations like that of FimH tipping type 1 pili (*20*). **Class 1-2** mAbs had varied binding affinity to *E. coli* FimG_nte_H, with K_on_ rates lower than binding to *E. coli* FimH_LD_ (K_on_ rates between 2×10^4^ to 1×10^5^ M^-1^s^-1^) (Fig. 3C, 3D, S6-7). This finding indicated that **Class 1-2** mAbs had a binding preference for the relaxed FimH conformation likely due to FimH_LD_ being the immunizing antigen.

**Fig. 3.**
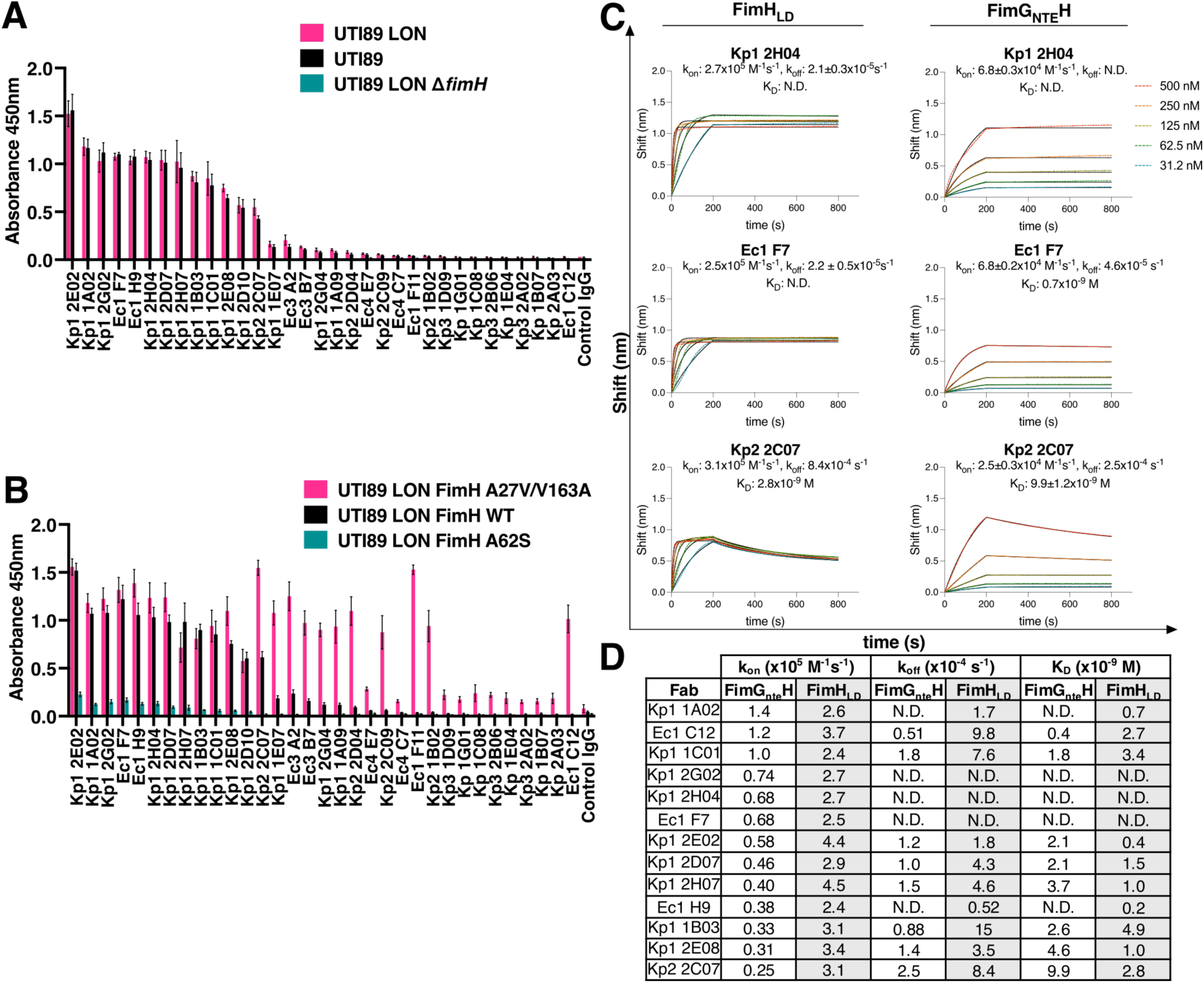
mAbs bind the relaxed conformation of FimH. (**A**) FimH mAb binding to UTI89 bacteria and (**B**) UTI89 overexpressing conformationally shifted FimH variants (n=3, error bars represent SEM). (**C**) Representative binding curves of Kp1 2H04, Ec1 F7, and Kp2 2C07 Fab binding FimH_LD_ and FimG_NTE_H. Results are from kinetic measurements of dilution series of one experiment. (**D**) Observed BLI binding kinetics to Ec FimG_NTE_H and Ec FimH_LD_. N.D. means not determined due to the off rate being below the detection limit.

### FimH_LD_ mAbs inhibit FimH mannosylated protein binding

The structural analysis showed mAbs recognizing epitopes near the mannose binding pocket suggesting mAbs may interfere with FimH binding. Thus, FimH_LD_ mAbs were tested for their ability to block *E. coli* and *K. pneumoniae* FimH_LD_ binding to highly mannosylated glycoprotein bovine submaxillary mucin (BSM). At a 5:1 molar ratio of mAb to FimH_LD_, assays measuring mAbs that inhibit binding to BSM in ELISA assays ranged from no inhibition to 85% inhibition compared to untreated control (Figure 4A). Only eight mAbs (representing epitope **Classes 1**-**3**) inhibited both FimH_LD_ proteins at greater than 50% at a 5:1 molar ratio. We selected mAbs from **Classes 1-3** from both *E. coli* and *K. pneumoniae* antigens with the highest inhibition to FimH_LD_ and the strongest binding to FimH tipping type 1 pili on the surface of bacteria to test for the ability to inhibit mannose-dependent *E. coli* bacterial hemagglutination of guinea pig erythrocytes. When present in high concentrations (17 uM), we found all mAbs tested can inhibit hemagglutination with more potency than ɑ-D-mannopyranoside. However, the FimH mAbs displayed greatly variable inhibition potency with 50% inhibition concentration ranging from 700 nM to above 17 uM (Fig. 4B). We selected Kp1 2H04 to further test for ability to block FimH_LD_ binding to mouse bladder tissue, as Kp1 2H04 had high inhibition in the FimH BSM ELISA and *E. coli* hemagglutination. At a 10:1 molar ratio of mAb to FimH_LD_ protein, Ec FimH_LD_ and Kp FimH_LD_ mixed with control IgG bound strongly to the bladder epithelial cells. However, Kp1 2H04 mAb treatment completely blocked Ec FimH_LD_ and Kp FimH_LD_ binding to mouse bladder tissue (Fig. 4C).

**Fig. 4.**
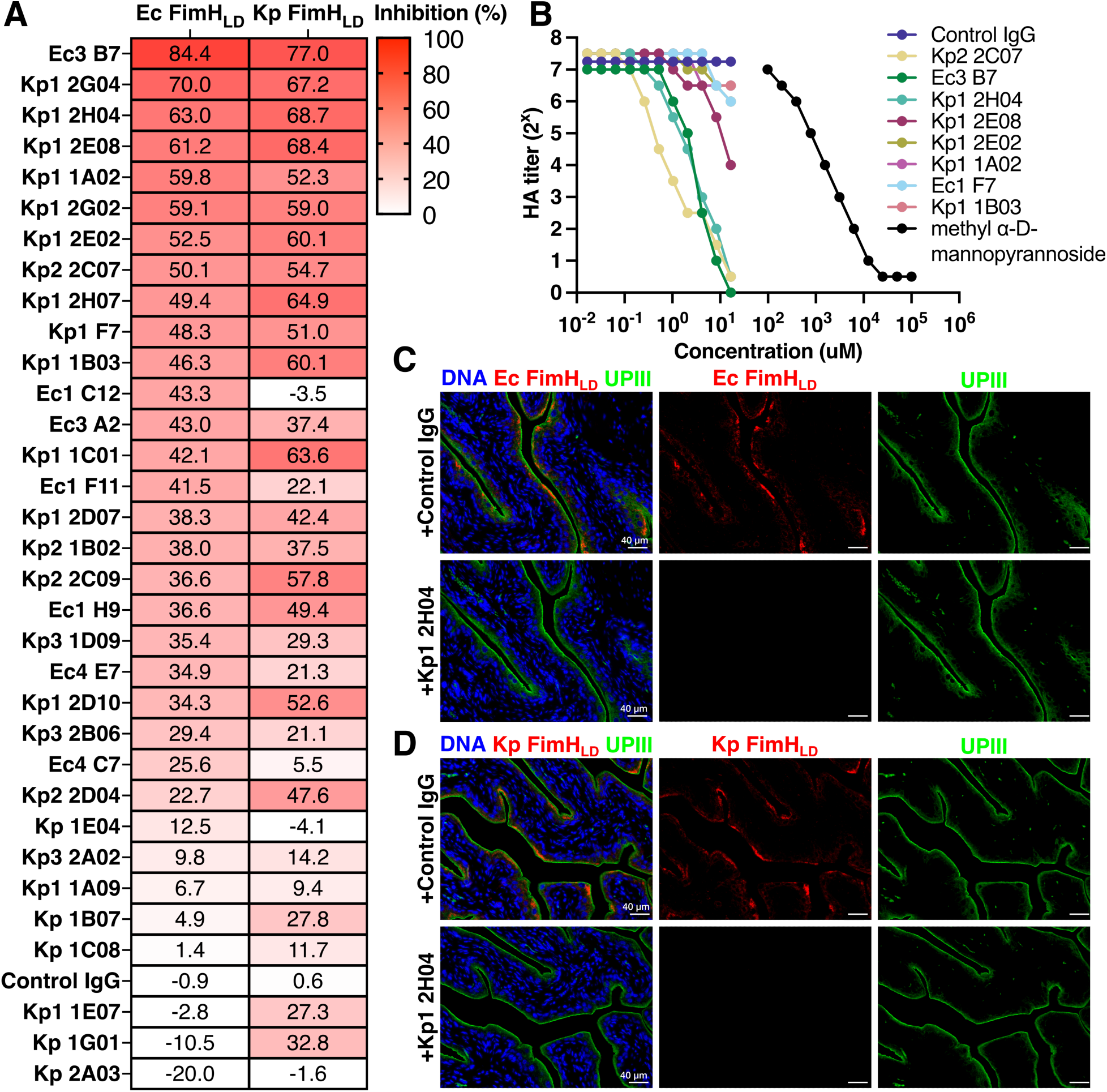
Relaxed-preferred mAbs can inhibit FimH binding. (**A**) Inhibition of FimH_LD_ binding to BSM at a 5:1 molar ratio of mAb to protein (n≥3). (**B**) Inhibition of UTI89 guinea pig erythrocyte hemagglutination (n≥2). (**C**) mAb inhibition of FimH_LD_ binding to C3H/HeN mouse bladders. mAb was pre-incubated with FimH_LD_ at a 10:1 molar ratio. Sections were stained with DNA dye Hoechst (blue), Ec or Kp FimH_LD_ (red) and antibody to uroplakin III (green). n=4-5 bladder sections with n=2 technical replicates.

### FimH_LD_ mAbs protect against UTI

To determine the ability of the mAbs to prevent UTI, we screened eight mAbs in a prophylactic model. We chose **Class 1-3** mAbs Ec1 F7, Kp1 2H04, Kp1 1A02, Kp1 2E02, Kp1 1B03, Kp1 2E08 mAbs, Kp2 2C07, and Ec3 B7 mAb for these assays since they bind with high-affinity to tip-like FimH, inhibit bacterial *E. coli* FimH binding, and represent three epitope classes from *E. coli* and *K. pneumoniae* FimH antigens. Each mAb was administered via intraperitoneal injection at 0.5 mg per mouse 24 hours before infection with *E. coli* UTI89. Bladder and kidney titers were enumerated 24 hours post-infection (Fig. 5A). There was no detectable IgG in urine before infection (24 hours after IP injection of mAbs) but mAbs were detected in the bladder approximately 3 to 6 hours after infection, consistent with bladder damage being necessary for antibodies to reach the urine as previously suggested (Fig. S8) (*30*). Three of the mAbs from epitope **Class 1**, (Ec1 F7, Kp1 2H04, and Kp1 1A02), resulted in significantly decreased bladder titers (∼1 log; P<0.05) and two **Class 1** mAbs (Ec1 F7 and Kp1 2H04) also significantly decreased kidney titers (∼0.5 log; P<0.05) (Fig. 5B and 5C). The effect of prophylactic administration of Kp1 2H04, one of the strongest inhibitors of bladder titers at 24 hpi, on formation of intracellular bacterial communities (IBCs) at 6 hpi was assessed. Kp1 2H04 significantly decreased the amount of bladder IBCs compared to the control IgG at 6 hpi (Fig. 5D-F).

**Fig. 5.**
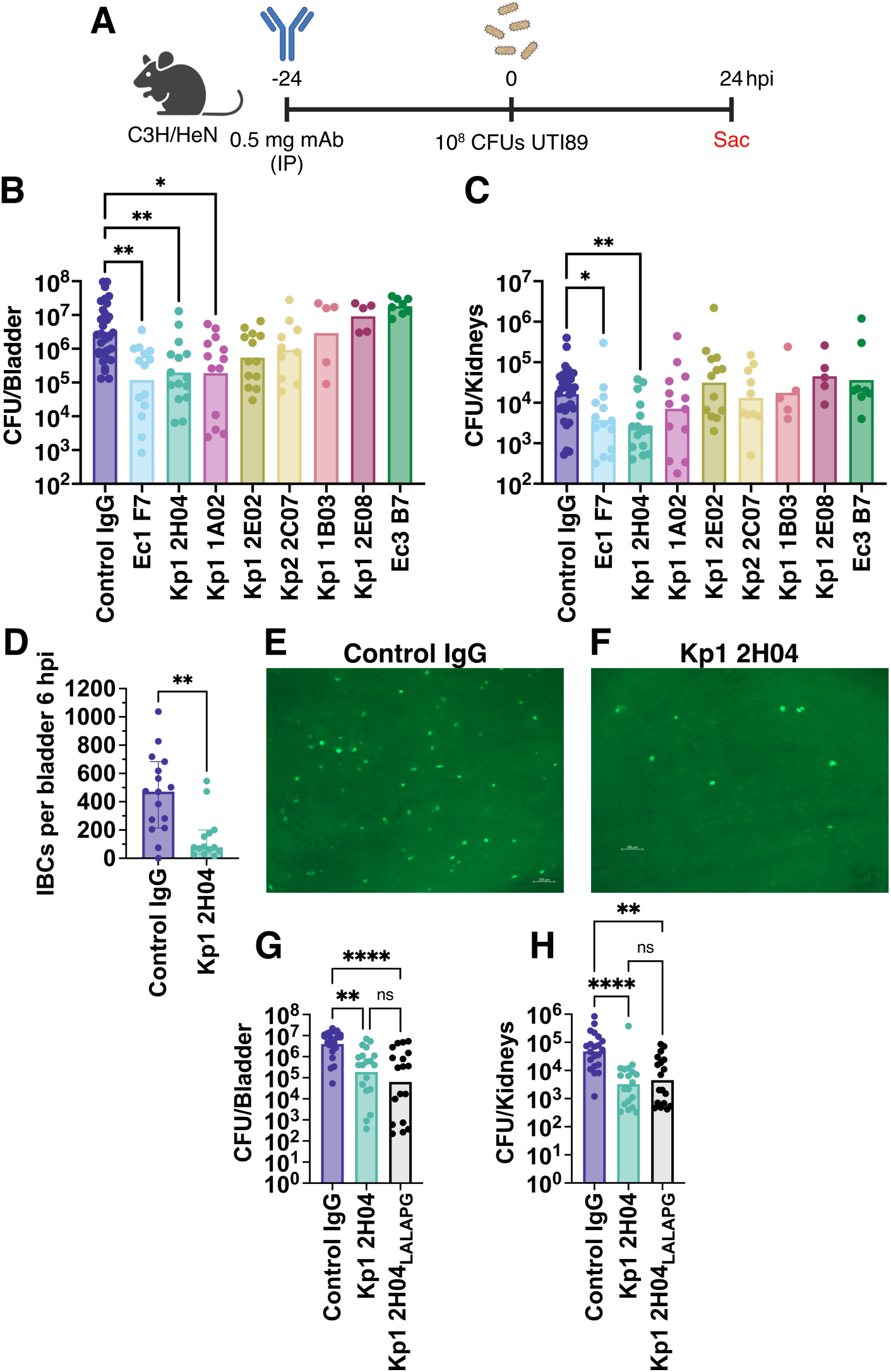
FimH mAbs protect from *E. coli* UTI. (**A**) 6-7 week old C3H/HeN mice were pretreated with 0.5 mg of mAb 24 h before infection with UTI89. (**B**) 24 hpi bladder and (**C**) kidney titers (For 1B03 and 2E08 n=5 with 1 independent replicate, for B7 n=8 with 1 independent replicate, for 2C07 n=10 with 2 independent replicates, for control IgG, F7, 2H04, 1A02, 2E02 n=13-33 with three independent replicates). (**D**) IBC counts at 6 hpi (n=16 for control IgG, n=14 for Kp1 2H04 with two independent replicates). Representative 5x magnification images of IBCs (green) in splayed mouse bladders for (**E**) control IgG and (**F**) Kp1 2H04 treatments. Scale bar represents 200 µm. 24 hpi bladder (**G**) and kidney titers (**H**) from the prophylactic model testing control IgG, 2H04, and 2H04_LALAPG_ (n=21 for control group, n=20 for 2H04, and n=19 for 2H04_LALAPG_ with three independent replicates). For **D**, a Mann-Whitney U test was used to evaluate statistical significance. For **B**, **C**, **G**, and **H** statistical comparisons were made using Kruskal-Wallis test (nonparametric ANOVA) with Dunn’s comparisons to the control group correcting for multiple comparisons. * P≤0.05, **P≤0.01, *** P≤0.0001., ****P≤0.00001.

To test if immune system signaling functions associated with the fragment crystallizable (Fc) region of mAbs were needed for protection, we created a LALAPG Fc variant of Kp1 2H04 (2H04_LALAPG_) which inhibits the ability of the mAb to bind Fc receptors or fix complement (*31*). 2H04_LALAPG_ had a comparable binding affinity to FimH_LD_ as 2H04 when assayed by ELISA (Fig. S9). When tested in the prophylactic model, 2H04_LALAPG_ inhibited UTI89 infection to the same degree as Kp1 2H04 in the bladder and kidney (Fig. 5G and Fig. 5H), suggesting that Kp1 2H04 prevents infection primarily by directly inhibiting FimH rather than through Fc receptor functions. To further test this, we repeated the prophylactic model but monitored the infection over 14 days. Kp1 2H04 treated mice resulted in lower urine and bladder titers over the 14 days. Still, this difference did not increase over time despite detectable amounts of mAb in the serum and bladder homogenates up to 14 days post-infection (Fig. S10) suggesting that the primary mechanism of the mAb is blocking initial FimH attachment to the bladder epithelium.

## Discussion

Carbapenem-resistant *Enterobacteriaceae* (CRE) and extended-spectrum beta-lactamase (ESBL) producing *Enterobacteriaceae* are listed as urgent and serious threats by the Centers for Diseases Control and Prevention (CDC) as they are resistant to numerous antibiotics needed to treat common infections, including UTIs (*32*, *33*). New antibiotic-sparing strategies are needed to treat these and other *Enterobacteriaceae* infections. mAbs have been exceptional drugs in treating cancer and viral infections (*34–38*) but few mAb therapies have been notably developed for treating or preventing bacterial infections. Here, we generated mAbs to neutralize FimH, the critical adhesin used by UPEC and *K. pneumoniae* to mediate UTI pathogenesis. We: **i)** generated mAbs that define four classes of anti-FimH mAbs that bind to distinct FimH epitopes; **ii)** identified cross-reactive mAbs with high affinity to *E. coli* and *K. pneumoniae* FimH; and **iii)** characterized the most potent FimH blocking antibodies in a structural and functional analysis. When we tested these mAbs *in vivo*, we found significant quantities of antibody could be detected in the urine of mice after (but not prior to) infection, suggesting that mAbs penetrate the urinary tract in the context of infection. Two **Class 1** Kp and Ec mAbs (Kp1 2H04 and Ec1 F7) significantly reduced bacterial titers in the bladder and kidneys when administered before UPEC infection in a robust mouse model of UTI through direct inhibition of FimH function. This is in contrast to previous studies suggesting that mAbs to the relaxed FimH conformation may stabilize a high-affinity conformation thereby increasing affinity to the bladder (*39*, *40*). These data suggest that some mAbs, which bind outside the binding pocket to a high-affinity relaxed-conformation by our structural analysis, inhibit FimH binding through steric hindrance preventing access to the binding pocket.

In the bladder, FimH binds uroplakin Ia, which forms an oligomeric complex with other uroplakin proteins in a dense crystalline lipid superstructure burying the mannose glycan deep in the complex (*41*). The structure of uroplakin plaques requires FimH to reach into a tight spatial pocket to bind mannose which may be sterically hindered by the binding of mAbs (Fig. S11). However, this proposed mechanism does not preclude the additional possibility of stabilizing the high-affinity conformation as shifting the conformational equilibrium towards the high-affinity state via mutations can also result in virulence attenuation (*20*, *42*, *43*). Bacteria with low-affinity tense-shifted FimH alleles may be able to evade vaccination or mAb therapy targeting the relaxed FimH conformation; however, bacteria with tense-shifted FimH alleles are attenuated in infection due to decreased ability to bind to the bladder epithelium (*21*, *42*, *43*). Nevertheless, the highly variable ability of antibodies to bind to tense or relaxed states of FimH suggests that the conformational equilibrium may provide an advantage *in vivo* by helping UPEC avoid antibodies that effectively bind to only one of the two states. Notably, we did not identify a mAb that directly binds to the FimH mannose-binding pocket, suggesting that vaccination of FimH in the relaxed state may function by providing antibodies that sterically inhibit FimH rather than directly blocking the mannose-binding pocket. Future studies are needed to determine the antigenicity of a tense state FimH and if tense state-specific anti-FimH mAbs are effective at protecting from UTI.

Our results show that FimH-inhibiting mAbs have promising antibiotic-sparing therapeutic potential to treat both UPEC and *K. pneumoniae* UTIs, lay the groundwork for identifying FimH mAbs with increased efficacy, and provide a roadmap to leverage mAbs to inhibit other bacterial uropathogenic CUP adhesins. While there have been few mAbs developed to treat bacterial infection, mAbs directed at treating UTI could offer unique advantages over antibiotics since they both avoid selection of antibiotic resistance and would have a sustained period of effectiveness. These mAbs may be deployed effectively in patients with highly recurrent UTI, who are often administered prophylactic antibiotics, or in hospital settings with high-risk UTI patients, such as patient populations requiring catheterization, in which UPEC and *K. pneumoniae* can repeatedly colonize catheters over many months (*44*).

## Acknowledgments

We acknowledge Amelia Coyne, Kevin Tamadonfar, Lily Liu, Maxwell Zimmerman, Nada Abdalla, Tingting Lei, and Wandy Beatty for their assistance during this study. The mouse timeline graphics in Figure 5 were created using BioRender.com. Cryo-EM microscopy and data collection were performed in part through the use of the Washington University Center for Cellular Imaging (WUCCI).

## Funding

National Institutes of Health grant R01AI029549 (SJH)

National Institutes of Health grant R37AI048689 (SJH)

National Institutes of Health grant U19AI157797 (SJH, AHE)

National Institutes of Health grant R01AI165915 (ALK)

Doris Duke Charitable Foundation 2019083 (ALK)

National Institutes of Health grant T32GM007067 (JMSB)

The Children’s Discovery Institute of Washington University and St. Louis Children’s Hospital grants CDI-CORE-2015-505

The Children’s Discovery Institute of Washington University and St. Louis Children’s Hospital grants CDI-CORE-2019-813

## Author contributions

Conceptualization: EDBL, JMSB, KWD, AHE, ALK, SJH

Methodology: EDBL, JMSB, DAS, SKM, PNA, MRT, IFF, AJS, JST, SMS, LO, NCG, JSP, KWD, AHE, ALK, SJH

Investigation: EDBL, JMSB, DAS, SKM, PNA, MRT, IFF, AJS, JST, SMS, LO, NCG, JSP, KWD

Resources: MRT, IFF, AJS, JST, SMS, LO, NCG, JSP, AHE, ALK, SJH

Formal analysis: EDBL, JMSB, DAS, SKM, PNA, MRT, IFF, AJS, JST, SMS, LO, NCG, JSP, KWD, AHE, ALK, SJH

Visualization: EDBL, JMSB, DAS, KWD, AHE, ALK, SJH

Funding acquisition: AHE, ALK, SJH

Project administration: AHE, ALK, SJH

Supervision: EDBL, JMSB, KWD, AHE, ALK, SJH

Writing – original draft: EDBL, JMSB, KWD, AHE, ALK, SJH

Writing – review & editing: EDBL, JMSB, DAS, SKM, PNA, MRT, IFF, AJS, JST, SMS, LO, NCG, JSP, KWD, AHE, ALK, SJH

## Competing interests

S.J.H. is on the advisory board of Sequoia Vaccines, Inc. S.J.H is a co-founder of Fimbrion Therapeutics that is developing FimH targeted therapies and may financially benefit if the company is successful. The Ellebedy laboratory received funding from Emergent BioSolutions, AbbVie, and Moderna that are unrelated to the data presented in the current study. A.H.E. has received consulting and speaking fees from InBios International, Fimbrion Therapeutics, RGAX, Mubadala Investment Company, Moderna, Pfizer, GSK, Danaher, Third Rock Ventures, Goldman Sachs and Morgan Stanley and is the founder of ImmuneBio Consulting. A. J. S., J.S.T., and A.H.E. are recipients of a licensing agreement with Abbvie that is unrelated to the data presented in the current study. All other authors declare that they have no competing interests.

## Data and materials availability

Cryo-EM data and protein models are deposited in PBD and EMDB with identification as follows: PDB 9CCS and EMDB EMD-45456 (2H04-FimH), PDB 9CCT and EMDB EMD-45457 (B7-FimH), PDB 9CCU and EMDB EMD-45458 (F7-FimH), and PDB 9CCW and EMDB EMD-45460 (2C07-FimH). All other data are available in the main text or the supplementary materials.

## References and Notes

1. X. Yang, H. Chen, Y. Zheng, S. Qu, H. Wang, F. Yi, Disease burden and long-term trends of urinary tract infections: A worldwide report. Front Public Health 10, 888205 (2022).

2. Z. Zeng, J. Zhan, K. Zhang, H. Chen, S. Cheng, Global, regional, and national burden of urinary tract infections from 1990 to 2019: an analysis of the global burden of disease study 2019. World J Urol 40, 755–763 (2022).

3. J. E. Simmering, F. Tang, J. E. Cavanaugh, L. A. Polgreen, P. M. Polgreen, The Increase in Hospitalizations for Urinary Tract Infections and the Associated Costs in the United States, 1998–2011. Open Forum Infect Dis 4, ofw281 (2017).

4. S. D. H. TM, R. PL, S. AE, G. K, S. WE, Risk factors for recurrent urinary tract infection in young women. J Infect Dis 182, 1177–1182 (2000).

5. C. W. Seymour, F. Gesten, H. C. Prescott, M. E. Friedrich, T. J. Iwashyna, G. S. Phillips, S. Lemeshow, T. Osborn, K. M. Terry, M. M. Levy, Time to Treatment and Mortality during Mandated Emergency Care for Sepsis. N Engl J Med 376, 2235–2244 (2017).

6. J. Hrbacek, P. Cermak, R. Zachoval, Current Antibiotic Resistance Trends of Uropathogens in Central Europe: Survey from a Tertiary Hospital Urology Department 2011-2019. Antibiotics (Basel) 9, 1–11 (2020).

7. B. Kot, Antibiotic Resistance Among Uropathogenic Escherichia coli. Pol J Microbiol 68, 403–415 (2019).

8. S. Navon-Venezia, K. Kondratyeva, A. Carattoli, Klebsiella pneumoniae: a major worldwide source and shuttle for antibiotic resistance. FEMS Microbiol Rev 41, 252–275 (2017).

9. A. L. Flores-Mireles, J. N. Walker, M. Caparon, S. J. Hultgren, Urinary tract infections: epidemiology, mechanisms of infection and treatment options. Nat Rev Microbiol 13, 269–284 (2015).

10. C. J. Murray, K. S. Ikuta, F. Sharara, L. Swetschinski, G. Robles Aguilar, A. Gray, C. Han, C. Bisignano, P. Rao, E. Wool, S. C. Johnson, A. J. Browne, M. G. Chipeta, F. Fell, S. Hackett, G. Haines-Woodhouse, B. H. Kashef Hamadani, E. A. P. Kumaran, B. McManigal, R. Agarwal, S. Akech, S. Albertson, J. Amuasi, J. Andrews, A. Aravkin, E. Ashley, F. Bailey, S. Baker, B. Basnyat, A. Bekker, R. Bender, A. Bethou, J. Bielicki, S. Boonkasidecha, J. Bukosia, C. Carvalheiro, C. Castañeda-Orjuela, V. Chansamouth, S. Chaurasia, S. Chiurchiù, F. Chowdhury, A. J. Cook, B. Cooper, T. R. Cressey, E. Criollo-Mora, M. Cunningham, S. Darboe, N. P. J. Day, M. De Luca, K. Dokova, A. Dramowski, S. J. Dunachie, T. Eckmanns, D. Eibach, A. Emami, N. Feasey, N. Fisher-Pearson, K. Forrest, D. Garrett, P. Gastmeier, A. Z. Giref, R. C. Greer, V. Gupta, S. Haller, A. Haselbeck, S. I. Hay, M. Holm, S. Hopkins, K. C. Iregbu, J. Jacobs, D. Jarovsky, F. Javanmardi, M. Khorana, N. Kissoon, E. Kobeissi, T. Kostyanev, F. Krapp, R. Krumkamp, A. Kumar, H. H. Kyu, C. Lim, D. Limmathurotsakul, M. J. Loftus, M. Lunn, J. Ma, N. Mturi, T. Munera-Huertas, P. Musicha, M. M. Mussi-Pinhata, T. Nakamura, R. Nanavati, S. Nangia, P. Newton, C. Ngoun, A. Novotney, D. Nwakanma, C. W. Obiero, A. Olivas-Martinez, P. Olliaro, E. Ooko, E. Ortiz-Brizuela, A. Y. Peleg, C. Perrone, N. Plakkal, A. Ponce-de-Leon, M. Raad, T. Ramdin, A. Riddell, T. Roberts, J. V. Robotham, A. Roca, K. E. Rudd, N. Russell, J. Schnall, J. A. G. Scott, M. Shivamallappa, J. Sifuentes-Osornio, N. Steenkeste, A. J. Stewardson, T. Stoeva, N. Tasak, A. Thaiprakong, G. Thwaites, C. Turner, P. Turner, H. R. van Doorn, S. Velaphi, A. Vongpradith, H. Vu, T. Walsh, S. Waner, T. Wangrangsimakul, T. Wozniak, P. Zheng, B. Sartorius, A. D. Lopez, A. Stergachis, C. Moore, C. Dolecek, M. Naghavi, Global burden of bacterial antimicrobial resistance in 2019: a systematic analysis. Lancet 399, 629–655 (2022).

11. J. J. Martinez, M. A. Mulvey, J. D. Schilling, J. S. Pinkner, S. J. Hultgren, Type 1 pilus-mediated bacterial invasion of bladder epithelial cells. EMBO J 19, 2803–2812 (2000).

12. D. A. Rosen, J. S. Pinkner, J. N. Walker, J. S. Elam, J. M. Jones, S. J. Hultgren, Molecular variations in Klebsiella pneumoniae and Escherichia coli FimH affect function and pathogenesis in the urinary tract. Infect Immun 76, 3346–3356 (2008).

13. L. K. McLellan, M. R. McAllaster, A. S. Kim, Ľ. Tóthová, P. D. Olson, J. S. Pinkner, A. L. Daugherty, T. N. Hreha, J. W. Janetka, D. H. Fremont, S. J. Hultgren, H. W. Virgin, D. A. Hunstad, A host receptor enables type 1 pilus-mediated pathogenesis of Escherichia coli pyelonephritis. PLoS Pathog 17 (2021).

14. M. A. Mulvey, J. D. Schilling, S. J. Hultgren, Establishment of a persistent Escherichia coli reservoir during the acute phase of a bladder infection. Infect Immun 69, 4572–4579 (2001).

15. D. A. Rosen, J. S. Pinkner, J. M. Jones, J. N. Walker, S. Clegg, S. J. Hultgren, Utilization of an intracellular bacterial community pathway in Klebsiella pneumoniae urinary tract infection and the effects of FimK on type 1 pilus expression. Infect Immun 76, 3337– 3345 (2008).

16. G. G. Anderson, J. J. Palermo, J. D. Schilling, R. Roth, J. Heuser, S. J. Hultgren, Intracellular bacterial biofilm-like pods in urinary tract infections. Science 301, 105–107 (2003).

17. V. P. O’Brien, T. J. Hannan, L. Yu, J. Livny, E. D. O. Roberson, D. J. Schwartz, S. Souza, C. L. Mendelsohn, M. Colonna, A. L. Lewis, S. J. Hultgren, A mucosal imprint left by prior Escherichia coli bladder infection sensitizes to recurrent disease. Nat Microbiol 2 (2016).

18. S. K. Russell, J. K. Harrison, B. S. Olson, H. J. Lee, V. P. O’Brien, X. Xing, J. Livny, L. Yu, E. D. O. Roberson, R. Bomjan, C. Fan, M. Sha, S. Estfanous, A. O. Amer, M. Colonna, T. S. Stappenbeck, T. Wang, T. J. Hannan, S. J. Hultgren, Uropathogenic Escherichia coli infection-induced epithelial trained immunity impacts urinary tract disease outcome. Nat Microbiol 8, 875–888 (2023).

19. D. Choudhury, A. Thompson, V. Stojanoff, S. Langermann, J. Pinkner, S. J. Hultgren, S. D. Knight, X-ray structure of the FimC-FimH chaperone-adhesin complex from uropathogenic Escherichia coli. Science 285, 1061–1066 (1999).

20. V. Kalas, J. S. Pinkner, T. J. Hannan, M. E. Hibbing, K. W. Dodson, A. S. Holehouse, H. Zhang, N. H. Tolia, M. L. Gross, R. V. Pappu, J. Janetka, S. J. Hultgren, Evolutionary fine-tuning of conformational ensembles in FimH during host-pathogen interactions. Sci Adv 3 (2017).

21. E. D. B. Lopatto, J. S. Pinkner, D. A. Sanick, R. F. Potter, L. X. Liu, J. Bazán Villicaña, K. O. Tamadonfar, Y. Ye, M. I. Zimmerman, N. C. Gualberto, K. W. Dodson, J. W. Janetka, D. A. Hunstad, S. J. Hultgren, Conformational ensembles in Klebsiella pneumoniae FimH impact uropathogenesis. Proc Natl Acad Sci U S A 121 (2024).

22. G. R. Eldridge, H. Hughey, L. Rosenberger, S. M. Martin, A. M. Shapiro, E. D’Antonio, K. G. Krejci, N. Shore, J. Peterson, A. S. Lukes, C. M. Starks, Safety and immunogenicity of an adjuvanted Escherichia coli adhesin vaccine in healthy women with and without histories of recurrent urinary tract infections: results from a first-in-human phase 1 study. Hum Vaccin Immunother 17, 1262–1270 (2021).

23. E. Perer, H. Stacey, T. Eichorn, H. Hughey, J. Lawrence, E. Cunningham, M. O. Johnson, K. Bacon, A. Kau, S. J. Hultgren, T. M. Hooton, J. L. Harris, Case report: Long-term follow-up of patients who received a FimCH vaccine for prevention of recurrent urinary tract infections caused by antibiotic resistant Enterobacteriaceae: a case report series. Front Immunol 15, 1359738 (2024).

24. S. Langermann, S. Palaszynski, M. Barnhart, G. Auguste, J. S. Pinkner, J. Burlein, P. Barren, S. Koenig, S. Leath, C. H. Jones, S. J. Hultgren, Prevention of mucosal Escherichia coli infection by FimH-adhesin-based systemic vaccination. Science 276, 607–611 (1997).

25. S. Langermann, R. Möllby, J. E. Burlein, S. R. Palaszynski, C. G. Auguste, A. DeFusco, R. Strouse, M. A. Schenerman, S. J. Hultgren, J. S. Pinkner, J. Winberg, L. Guldevall, M. Söderhäll, K. Ishikawa, S. Normark, S. Koenig, Vaccination with FimH adhesin protects cynomolgus monkeys from colonization and infection by uropathogenic Escherichia coli. J Infect Dis 181, 774–778 (2000).

26. L. Von Boehmer, C. Liu, S. Ackerman, A. D. Gitlin, Q. Wang, A. Gazumyan, M. C. Nussenzweig, Sequencing and cloning of antigen-specific antibodies from mouse memory B cells. Nature Protocols 2016 11:10 11, 1908–1923 (2016).

27. M. S. Conover, S. Ruer, J. Taganna, V. Kalas, H. De Greve, J. S. Pinkner, K. W. Dodson, H. Remaut, S. J. Hultgren, Inflammation-Induced Adhesin-Receptor Interaction Provides a Fitness Advantage to Uropathogenic E. coli during Chronic Infection. Cell Host Microbe 20, 482–492 (2016).

28. D. I. Kisiela, P. Magala, G. Interlandi, L. A. Carlucci, A. Ramos, V. Tchesnokova, B. Basanta, V. Yarov-Yarovoy, H. Avagyan, A. Hovhannisyan, W. E. Thomas, R. E. Stenkamp, R. E. Klevit, E. V. Sokurenko, Toggle switch residues control allosteric transitions in bacterial adhesins by participating in a concerted repacking of the protein core. PLoS Pathog 17 (2021).

29. I. Le Trong, P. Aprikian, B. A. Kidd, M. Forero-Shelton, V. Tchesnokova, P. Rajagopal, V. Rodriguez, G. Interlandi, R. Klevit, V. Vogel, R. E. Stenkamp, E. V. Sokurenko, W. E. Thomas, Structural basis for mechanical force regulation of the adhesin FimH via finger trap-like beta sheet twisting. Cell 141, 645–655 (2010).

30. A. L. Flores-Mireles, J. N. Walker, A. Potretzke, H. L. Schreiber, J. S. Pinkner, T. M. Bauman, A. M. Park, A. Desai, S. J. Hultgren, M. G. Caparon, Antibody-Based Therapy for Enterococcal Catheter-Associated Urinary Tract Infections. mBio 7 (2016).

31. T. Schlothauer, S. Herter, C. F. Koller, S. Grau-Richards, V. Steinhart, C. Spick, M. Kubbies, C. Klein, P. Umaña, E. Mössner, Novel human IgG1 and IgG4 Fc-engineered antibodies with completely abolished immune effector functions. Protein Eng Des Sel 29, 457–466 (2016).

32. 2019 Antibiotic Resistance Threats Report | Antimicrobial Resistance | CDC. https://www.cdc.gov/antimicrobial-resistance/data-research/threats/index.html.

33. Antimicrobial Resistance Threats in the United States, 2021-2022 | Antimicrobial Resistance | CDC. https://www.cdc.gov/antimicrobial-resistance/data-research/threats/update-2022.html.

34. C. H. Chau, P. S. Steeg, W. D. Figg, Antibody–drug conjugates for cancer. The Lancet 394, 793–804 (2019).

35. D. Corti, L. A. Purcell, G. Snell, D. Veesler, Tackling COVID-19 with neutralizing monoclonal antibodies. Cell 184, 3086–3108 (2021).

36. M. P. Griffin, Y. Yuan, T. Takas, J. B. Domachowske, S. A. Madhi, P. Manzoni, E. A. F. Simões, M. T. Esser, A. A. Khan, F. Dubovsky, T. Villafana, J. P. DeVincenzo, Single-Dose Nirsevimab for Prevention of RSV in Preterm Infants. New England Journal of Medicine 383, 415–425 (2020).

37. S. Mulangu, L. E. Dodd, R. T. Davey, O. Tshiani Mbaya, M. Proschan, D. Mukadi, M. Lusakibanza Manzo, D. Nzolo, A. Tshomba Oloma, A. Ibanda, R. Ali, S. Coulibaly, A. C. Levine, R. Grais, J. Diaz, H. C. Lane, J.-J. Muyembe-Tamfum, A Randomized, Controlled Trial of Ebola Virus Disease Therapeutics. New England Journal of Medicine 381, 2293–2303 (2019).

38. J. J. Lim, A. C. Nilsson, M. Silverman, N. Assy, P. Kulkarni, J. M. McBride, R. Deng, C. Li, X. Yang, A. Nguyen, P. Horn, M. Maia, A. Castro, M. C. Peck, J. Galanter, T. Chu, E. M. Newton, J. A. Tavel, A phase 2 randomized, double-blind, placebo-controlled trial of MHAA4549A, a monoclonal antibody, plus oseltamivir in patients hospitalized with severe influenza a virus infection. Antimicrob Agents Chemother 64 (2020).

39. V. Tchesnokova, P. Aprikian, O. Yakovenko, C. LaRock, B. Kidd, V. Vogel, W. Thomas, E. Sokurenko, Integrin-like allosteric properties of the catch bond-forming FimH adhesin of Escherichia coli. J Biol Chem 283, 7823–7833 (2008).

40. V. Tchesnokova, P. Aprikian, D. Kisiela, S. Gowey, N. Korotkova, W. Thomas, E. Sokurenko, Type 1 fimbrial adhesin FimH elicits an immune response that enhances cell adhesion of Escherichia coli. Infect Immun 79, 3895–3904 (2011).

41. H. Yanagisawa, Y. Kita, T. Oda, M. Kikkawa, Cryo-EM elucidates the uroplakin complex structure within liquid-crystalline lipids in the porcine urothelial membrane. Commun Biol 6 (2023).

42. D. J. Schwartz, V. Kalas, J. S. Pinkner, S. L. Chen, C. N. Spaulding, K. W. Dodson, S. J. Hultgren, Positively selected FimH residues enhance virulence during urinary tract infection by altering FimH conformation. Proc Natl Acad Sci U S A 110, 15530–15537 (2013).

43. S. L. Chen, C. S. Hung, J. S. Pinkner, J. N. Walker, C. K. Cusumano, Z. Li, J. Bouckaert, J. I. Gordon, S. J. Hultgren, Positive selection identifies an in vivo role for FimH during urinary tract infection in addition to mannose binding. Proc Natl Acad Sci U S A 106, 22439–22444 (2009).

44. T. M. Nye, Z. Zou, C. L. P. Obernuefemann, J. S. Pinkner, E. Lowry, K. Kleinschmidt, K. Bergeron, A. Klim, K. W. Dodson, A. L. Flores-Mireles, J. N. Walker, D. G. Wong, A. Desai, M. G. Caparon, S. J. Hultgren, Microbial co-occurrences on catheters from long-term catheterized patients. Nature Communications 2024 15:1 15, 1–13 (2024).

45. J. Bouckaert, J. Berglund, M. Schembri, E. De Genst, L. Cools, M. Wuhrer, C. S. Hung, J. Pinkner, R. Slättegård, A. Zavialov, D. Choudhury, S. Langermann, S. J. Hultgren, L. Wyns, P. Klemm, S. Oscarson, S. D. Knight, H. De Greve, Receptor binding studies disclose a novel class of high-affinity inhibitors of the Escherichia coli FimH adhesin. Mol Microbiol 55, 441–455 (2005).

46. R. A. Laskowski, Enhancing the functional annotation of PDB structures in PDBsum using key figures extracted from the literature. Bioinformatics 23, 1824–1827 (2007).

47. W. B. Alsoussi, J. S. Turner, J. B. Case, H. Zhao, A. J. Schmitz, J. Q. Zhou, R. E. Chen, T. Lei, A. A. Rizk, K. M. McIntire, E. S. Winkler, J. M. Fox, N. M. Kafai, L. B. Thackray, A. O. Hassan, F. Amanat, F. Krammer, C. T. Watson, S. H. Kleinstein, D. H. Fremont, M. S. Diamond, A. H. Ellebedy, A Potently Neutralizing Antibody Protects Mice against SARS-CoV-2 Infection. The Journal of Immunology 205, 915–922 (2020).

48. T. Tiller, C. E. Busse, H. Wardemann, Cloning and expression of murine Ig genes from single B cells. J Immunol Methods 350, 183–193 (2009).

49. M. Ehlers, H. Fukuyama, T. L. McGaha, A. Aderem, J. V. Ravetch, TLR9/MyD88 signaling is required for class switching to pathogenic IgG2a and 2b autoantibodies in SLE. J Exp Med 203, 553–561 (2006).

50. I. Y. Ho, J. J. Bunker, S. A. Erickson, K. E. Neu, M. Huang, M. Cortese, B. Pulendran, P. C. Wilson, Refined protocol for generating monoclonal antibodies from single human and murine B cells. J Immunol Methods 438, 67–70 (2016).

51. C. W. Davis, K. J. L. Jackson, A. K. McElroy, P. Halfmann, J. Huang, C. Chennareddy, A. E. Piper, Y. Leung, C. G. Albariño, I. Crozier, A. H. Ellebedy, J. Sidney, A. Sette, T. Yu, S. C. A. Nielsen, A. J. Goff, C. F. Spiropoulou, E. O. Saphire, G. Cavet, Y. Kawaoka, A. K. Mehta, P. J. Glass, S. D. Boyd, R. Ahmed, Longitudinal Analysis of the Human B Cell Response to Ebola Virus Infection. Cell 177, 1566–1582.e17 (2019).

52. S. J. Hultgren, W. R. Schwan, A. J. Schaeffer, J. L. Duncan, Regulation of production of type 1 pili among urinary tract isolates of Escherichia coli. Infect Immun 54, 613 (1986).

53. A. Punjani, J. L. Rubinstein, D. J. Fleet, M. A. Brubaker, cryoSPARC: algorithms for rapid unsupervised cryo-EM structure determination. Nat Methods 14, 290–296 (2017).

54. A. Punjani, H. Zhang, D. J. Fleet, Non-uniform refinement: adaptive regularization improves single-particle cryo-EM reconstruction. Nat Methods 17, 1214–1221 (2020).

55. T. Bepler, A. Morin, M. Rapp, J. Brasch, L. Shapiro, A. J. Noble, B. Berger, Positive-unlabeled convolutional neural networks for particle picking in cryo-electron micrographs. Nat Methods 16, 1153–1160 (2019).

56. A. Waterhouse, M. Bertoni, S. Bienert, G. Studer, G. Tauriello, R. Gumienny, F. T. Heer, T. A. P. De Beer, C. Rempfer, L. Bordoli, R. Lepore, T. Schwede, SWISS-MODEL: homology modelling of protein structures and complexes. Nucleic Acids Res 46, W296– W303 (2018).

57. E. F. Pettersen, T. D. Goddard, C. C. Huang, E. C. Meng, G. S. Couch, T. I. Croll, J. H. Morris, T. E. Ferrin, UCSF ChimeraX: Structure visualization for researchers, educators, and developers. Protein Sci 30, 70–82 (2021).

58. D. Liebschner, P. V. Afonine, M. L. Baker, G. Bunkoczi, V. B. Chen, T. I. Croll, B. Hintze, L. W. Hung, S. Jain, A. J. McCoy, N. W. Moriarty, R. D. Oeffner, B. K. Poon, M. G. Prisant, R. J. Read, J. S. Richardson, D. C. Richardson, M. D. Sammito, O. V. Sobolev, D. H. Stockwell, T. C. Terwilliger, A. G. Urzhumtsev, L. L. Videau, C. J. Williams, P. D. Adams, Macromolecular structure determination using X-rays, neutrons and electrons: recent developments in Phenix. Acta Crystallogr D Struct Biol 75, 861– 877 (2019).

59. P. Emsley, B. Lohkamp, W. G. Scott, K. Cowtan, Features and development of Coot. Acta Crystallogr D Biol Crystallogr 66, 486–501 (2010).

60. T. J. Hannan, D. A. Hunstad, A murine model for escherichia coli urinary tract infection. Methods in Molecular Biology 1333, 159–175 (2016).

61. B. Xie, G. Zhou, S. Y. Chan, E. Shapiro, X. P. Kong, X. R. Wu, T. T. Sun, C. E. Costello, Distinct glycan structures of uroplakins Ia and Ib: structural basis for the selective binding of FimH adhesin to uroplakin Ia. J Biol Chem 281, 14644–14653 (2006).

62. K. A. Datsenko, B. L. Wanner, One-step inactivation of chromosomal genes in Escherichia coli K-12 using PCR products. Proc Natl Acad Sci U S A 97, 6640–6645 (2000).

63. S. E. Greene, M. E. Hibbing, J. Janetka, S. L. Chen, S. J. Hultgren, Human urine decreases function and expression of type 1 pili in uropathogenic Escherichia coli. mBio 6 (2015).

64. M. Kostakioti, M. Hadjifrangiskou, C. K. Cusumano, T. J. Hannan, J. W. Janetka, S. J. Hultgren, Distinguishing the contribution of type 1 pili from that of other QseB-misregulated factors when QseC is absent during urinary tract infection. Infect Immun 80, 2826–2834 (2012).

65. K. J. Wright, P. C. Seed, S. J. Hultgren, Uropathogenic Escherichia coli flagella aid in efficient urinary tract colonization. Infect Immun 73, 7657–7668 (2005).

66. I. U. Mysorekar, S. J. Hultgren, Mechanisms of uropathogenic Escherichia coli persistence and eradication from the urinary tract. Proc Natl Acad Sci U S A 103, 14170– 14175 (2006).

67. I. Y. Ho, J. J. Bunker, S. A. Erickson, K. E. Neu, M. Huang, M. Cortese, B. Pulendran, P. C. Wilson, Refined protocol for generating monoclonal antibodies from single human and murine B cells. J Immunol Methods 438, 67–70 (2016).

