## Supplemental Materials for "Monoclonal antibodies targeting the FimH adhesin protect against uropathogenic *E. coli* UTI"

**The PDF file includes:**

Materials and Methods
Supplementary Text
Figs. S1 to S11
Tables S1 to S5

### Materials and Methods

Protein generation and purification

FimCH, $FimH_{LD}$ and FmlH lectin domain truncated proteins were purified as previously described (*21*, *27*, *42*, *45*). Briefly, proteins were expressed and isolated from crude periplasmic preparations using affinity chromatography. *In vitro* DSE using $FimG_{NTE}$ peptide was performed and purified as previously published (*20*). $FimH_{LD}$ labeled with EZ-Link$^{TM}$ NHS-PEG4 Biotinylation Kit (ThermoFisher) or Alexa Fluor 647 with NHS Ester conjugation (Invitrogen) was generated according to the manufacturer's instructions. Surface topology diagram of FimH was generated using PDBsum (*46*).

Mouse immunization for monoclonal antibody (mAb) generation

All procedures involving animals were performed in accordance with the guidelines of the Institutional Animal Care and Use Committee (IACUC) of Washington University in Saint Louis. Female C57BL/6J mice (Jackson Laboratories) were immunized intramuscularly with 30 μg *E. coli* $FimH_{LD}$ or 25 μg *K. pneumoniae* $FimH_{LD}$ emulsified with AddaVax (InvivoGen). Four weeks later, mice were boosted with a second dose of $FimH_{LD}$ emulsified with AddaVax. One control mouse received PBS emulsified with AddaVax according to the same schedule. Draining iliac and inguinal lymph nodes were harvested 5 days after the boost for plasmablast sorting.

Cell Sorting for mAb generation

Staining for sorting was performed using fresh lymph node single cell suspensions in PBS supplemented with 2% FBS and 1mM EDTA (P2). Cells were stained for 30 min on ice with CD138-BV421 (281-2, 1:200), CD4-PerCP (GK1.5, 1:100), CD19-PE (6D5, 1:200), B220-PE-D594 (RA3-6B2, 1:200), CD38-PE-Cy7 (90, 1:200), Fas-APC (SA367H8, 1:400), IgD-APC-Cy7 (11-26c.2a, 1:100), and Zombie Aqua (all Biolegend) diluted in P2. Cells were washed twice and single plasmablasts (B220lo CD138+ IgDlo CD19+ CD4- live singlet lymphocytes) were sorted using a FACSAria II into 96-well plates containing 2 μL Lysis Buffer (Clontech) supplemented with 1 U/μL RNase inhibitor (NEB) and immediately frozen on dry ice.

Monoclonal antibody (mAb) and Fragment antigen binding (Fab) generation

Antibodies were cloned as previously described (*26*, *47*). In brief, VH, Vκ, and Vλ genes were amplified by reverse transcriptase-polymerase chain reaction (RT-PCR) and nested PCR from singly-sorted plasmablasts using cocktails of primers specific for IgG, IgM/A, Igκ, and Igλ using first round and nested primer sets (*26*, *47–49*) (Table S3) and then sequenced. Clonally related cells were identified by the same length and composition of IGHV, IGHJ and heavy-chain CDR3 and shared somatic hypermutation at the nucleotide level. To generate recombinant antibodies, heavy chain V-D-J and light chain V-J fragments were PCR-amplified from 1st round PCR products with mouse variable gene forward primers and joining gene reverse primers having 5' extensions for cloning by Gibson assembly as previously described (*50*) (Table S3), and were cloned into pABVec6W antibody expression vectors (*51*) in frame with either human IgG, IgK, or IgL constant domain. Plasmids were co-transfected at a 1:2 heavy to light chain ratio into Expi293F cells using the Expifectamine 293 Expression Kit (Thermo Fisher), and antibodies were purified with protein A agarose (Invitrogen). For monovalent Fab generation, the VH segment of selected antibodies were cloned into a Fab expression vector with a thrombin cleavage site preceding a 6xHis tag by GenScript. Fab and light chain plasmids were co-transfected into Expi293F cells for expression and purified with HisPur Ni-NTA resin (Thermo Scientific). For controls in experiments, IgG mAb 2B04, specific for SARS-CoV-2 receptor binding domain (*47*) was used.

ELISAs

To generate ELISA binding curves to $FimH_{LD}$ and $FmlH_{LD}$ adhesin truncates, plates were first coated with the antigen (0.1 ug/mL) overnight. Plates were washed once with PBS supplemented with Tween-20 (PBS-T, 0.05%) and blocked for 2 h with PBS-T with 10% FBS. mAbs were serially diluted starting at 30 ug/mL and added to the plate to bind for 1h. Plates were washed 3x in PBS-T. mAb binding was detected with anti-human IgG (Jackson ImmunoResearch, 1:2,500 dilution) for 1 h before washing 3 times in PBS-T and developed with O-phenylenediamine dihydrochloride in citrate buffer (Sigma). Reactions were quenched with 1M HCl and absorbance was read at 490 nm.

To measure inhibition of $FimH_{LD}$ binding, plates were coated with BSM (10 ug/mL) overnight. Plates were blocked for 2 h with 1x PBS with 1% BSA. $FimH_{LD}$ (2.5 ug/mL) was mixed with a serial curve of mAb (to ensure dose-dependent inhibition) for 1 h. $FimH_{LD}$ mAb mixtures were then added to the plate to let bind for 1 h at RT. Plates were washed 3 times with PBS-T and incubated with anti-streptavidin-HRP (BD Pharmagen) for 1 h before washing 3 times with PBS-T and development with TMB substrate (BD Pharmagen). Reactions were quenched with 1M $H_2SO_4$ and absorbance at 450 nm was recorded.

For measurements of the reactivity of mAbs to bacteria, bacteria were grown statically 2x24 (grown for 24 h and subcultured 1:1000 for another 24h growth period) in LB and normalized to $OD_{600}$ = 1.0 in 1x PBS. Bacteria (100 uL) were added to the plate, spun down 5 min at 3000 rcf, and allowed to bind for 1 h. The supernatant was decanted and formalin (10%) was added to the wells to fix bacteria for 10 min. Plates were washed 3x in PBS-T and then assayed and developed with the same protocol as the ELISA binding curves to adhesin truncates.
To measure levels of humanized IgG (mAbs) in serum, urines and bladder homogenates, plates were coated with diluted 1:100 serum, 1:10 mouse urine, or 1:2 bladder homogenates along with a standard curve of hIgG isotype overnight at 4°C. Plates were then blocked for 2 hours at room temperature with 1x PBS with 1% BSA. Then, plates were washed three times with PBS-T. To measure hIgG levels, plates were then incubated with 1:10,000 dilution of goat anti-Human IgG H&L (HRP) preadsorbed IgG (Abcam, ab97175) for 1 hr at room temperature. For plate development, TMB substrate reagent was added and incubated for approximately 5 minutes at room temperature. Reactions were quenched with 3M HCl and absorbance at 450 nm was recorded.

Generation of FimH surface mutants

Surface FimH (J96) mutants were generated via one-step mutagenesis (19055817) using a pBAD33.1 vector plasmid encoding for FimH (J96 strain) template, Pfu Ultra HF polymerase (Agilent, NC9666083), and the primers listed in Table S3. After PCR amplification, reactions were digested with DpnI (NEB Biolabs, R0176S) to remove the original template from the reaction products. PCR reaction products were transformed into *E. coli* DH5ɑ for ligation. Plasmids confirmed by Sanger sequencing were then transformed into *Escherichia coli* C600 *Δfim* expression strain.

Epitope mapping

Epitope mapping of FimH mAbs to *E. coli* $FimC_{his}H$ was performed using a modified ELISA technique. *E. coli* strain C600 *Δfim* carrying pBAD33 plasmids encoding FimH mutants and a ptrc99a plasmid encoding $FimC_{his}$, were grown in LB to an OD of 0.6-0.8 and then induced with 0.1mM IPTG and 0.05% arabinose for 1 h. Cells were harvested and periplasm extracts containing $FimC_{his}H$ variants were obtained (*45*). $FimC_{his}H$ periplasmic extracts were titrated using anti-FimH sera to normalize the amount of FimH. Normalized $FimC_{his}H$ periplasm was used to coat plates for 1 h at room temperature. Plates were blocked for 2 h with 1x PBS with 1% BSA and mAbs (0.1 ug/mL, except for low-binding mAb 2A02 1ug/ml was used) were allowed to bind for 1 h. Plates were washed in 1x PBS-T before detection with anti-human IgG (Jackson ImmunoResearch, 1:1000 dilution) and developed with TMB substrate (BD Pharmagen) and $H_2SO_4$ as outlined above. mAb binding to each $FimC_{his}H$ mutant was normalized to WT $FimC_{his}H$ binding. Variants that decreased binding below 10% of WT binding and clustered together (3 or more residues) were considered an epitope. Mapping data was visualized and clustered (one-minus Pearson correlation) in Morpheus (https://software.broadinstitute.org/morpheus).

Western blotting

Western blotting to detect FimA in bacterial lysates was performed as previously published (*21*). Bacteria grown 2x24 statically was normalized to optical density at 600nm of 1.0 and were acid treated with HCl and boiled to disrupt FimA DSE interactions. To measure FimA, rabbit antitype 1 pili (1:2000) was used. A secondary antibody of goat anti-rabbit-HRP (1:10,000, KPL) was used to detect followed by development with SuperSignalTM West Femto Maximum Sensitivity Substrate (Thermo Fisher). Images were obtained on a BioRad ChemiDoc system. A colorimetric image (to view protein size ladder) was overlaid on the chemiluminescent image (detection signal) to create the figure reported in this study.

Biolayer interferometry (BLI) studies

Kinetic binding studies were performed on an Octet Red instrument (ForteBio). Avi-tag biotinylated antibody fragments (Fab) were loaded up to 2.5 nm onto Streptavidin sensor tips (Sartorius) that were pre-equilibrated in HEPES Buffered Saline (HBS) with 0.05% Tween-20 and 1% BSA (kinetic buffer A). Diluted antigens (Ec $FimH_{LD}$, Ec $FimG_{nte}H$) were monitored for 200s of association and 600s of dissociation in kinetic buffer A. Loaded sensor tips dipping in kinetic buffer A were used as reference sensors. Reference subtracted kinetic traces were used to calculate kinetic rate constants ($k_{on}$, $k_{off}$) and equilibrium dissociation rate constant ($K_D$) using a Langmuir 1:1 binding model. Resulting binding traces and fits were plotted with GraphPad Prism v10.

Hemagglutination inhibition (HAI) assay

*E. coli* guinea pig erythrocyte hemagglutination inhibition assays were performed as previously described (*52*). Briefly, mAbs were serially diluted in microtiter plates, and 25uL of bacterial suspension (serially diluted from $OD_{600}$=10.0 in 1x PBS) was added to each well. After incubation for 10 min at room temperature, 25 ul ($OD_{640}$=2.0) guinea pig erythrocytes in 1x PBS were added for a final volume of 50ul. The plates were incubated at 4°C overnight. For each mAb concentration, the HA titer was defined as the greatest dilution of bacteria that caused hemagglutination.

Cryo-EM data collection and analysis

Fab and FimCH protein were mixed at a ratio of 1.2:1 and dialyzed into 20mM HEPES pH 7.5 with 50mM NaCl. Complexes were flash frozen on EM grids in liquid ethane using an FEI Vitrobot (ThermoFisher) and imaged on Titan Krios (2H04 complex) or Glacios (F7, B7 and 2C07 complexes) microscopes using a Falcon 4 electron detector (Thermo Fisher). Movies were processed in Cryosparc v4.4.1 (*53*, *54*) and particles were picked using Topaz (*55*). Densities were post-processed using DeepEMhancer (*55*) for model building. The collection parameters and workflow are described in more detail in Fig. S3. Initial Fab models were built using homology models in SwissModel (*56*). $FimH_{LD}$ model was generated by trimming and threading the UTI89 FimH sequence on PDB 1KLF. Rough models were initially docked in ChimeraX (*57*) before multiple rounds of real space refinement in Phenix v1.20.1 (*58*) with manual editing in COOT v0.9.6 (*59*). Refinement statistics are shown in Table S4.

Immunofluorescence studies

7-8 week old female C3H/HeN mice (Envigo) bladders were fixed in formalin, embedded in paraffin, and sectioned. Tissue sections were heat deparaffinized and rehydrated in xylene, followed by stepwise hydration in 100% ethanol, 90% ethanol, 75% ethanol, 50% ethanol to 30% ethanol (each step having a 5 min incubation in fresh solution). Slides were rinsed in 1x PBS followed by blocking solution (1x PBS with 5% fetal bovine serum). Primary mouse antibody to uroplakin IIIa (Progen, 1:50) was allowed to bind overnight at 4°C. Slides were washed in 1x PBS and a secondary anti-mouse Alexa fluor 488 antibody (Invitrogen, 1:1000) was allowed to bind for 2 h, and then washed again. $FimH_{LD}$-Alexa Fluor 647 (588 nM) was mixed with mAb (6 uM) for 20 min at room temperature in 1x PBS. $FimH_{LD}$ mAb mixtures were applied to the section along with Hoechst DNA dye (8 uM) for 20 min at room temperature. The slides were washed again in 1x PBS and allowed to dry. ProLong$^{TM}$ gold antifade mountant (Invitrogen) was added and slides were imaged using the confocal function of a Zeiss Cell Observer Spinning Disk Confocal Microscope with a 10x air objective lens.

Splayed bladders were analyzed with a Zeiss Axio Observer D1 inverted fluorescence microscope equipped with an X-Cite120 mini LED light source (Excelitis Technologies) and DAPI, GFP, DsRed, and Cy5 filter sets. EC Plan-Neofluar (NA 0.075) 2.5X and EC Plan-Neofluar (NA 0.15) 5X objectives (Zeiss), an Axiocam 503 color camera (Zeiss), and ZEN 2 (blue version) software were used for image acquisition.

Mouse infection experiments

For acute and 2-week infection models, 7-8 week old female C3H/HeN mice (Envigo) were infected with $2x10^8$ CFUs of UTI89 as previously described (*60*). Intraperitoneal injections of mAb were given in 1x PBS buffer 24 h before infection. Urines were taken by clean catch at specified time points. For obtaining serum, 3 to 4 mice per treatment group were bled at specific time points during the length of the experiment via submental bleeding method. At the conclusion of the experimental time points, mice were humanely sacrificed and bladder and kidney organs were homogenized and tittered. For screening mAbs in the prophylactic infection model with UTI89, if a phenotype was observed after 1 replicate, the experiment was repeated an additional 1-2 times. All studies were approved and performed in accordance with the guidelines set by the Committee for Animal Studies at Washington University School of Medicine under IACUC protocols 21-0341 and 21-0394. (IACUC Protocol Approval Animal Welfare Assurance # D16-00245).

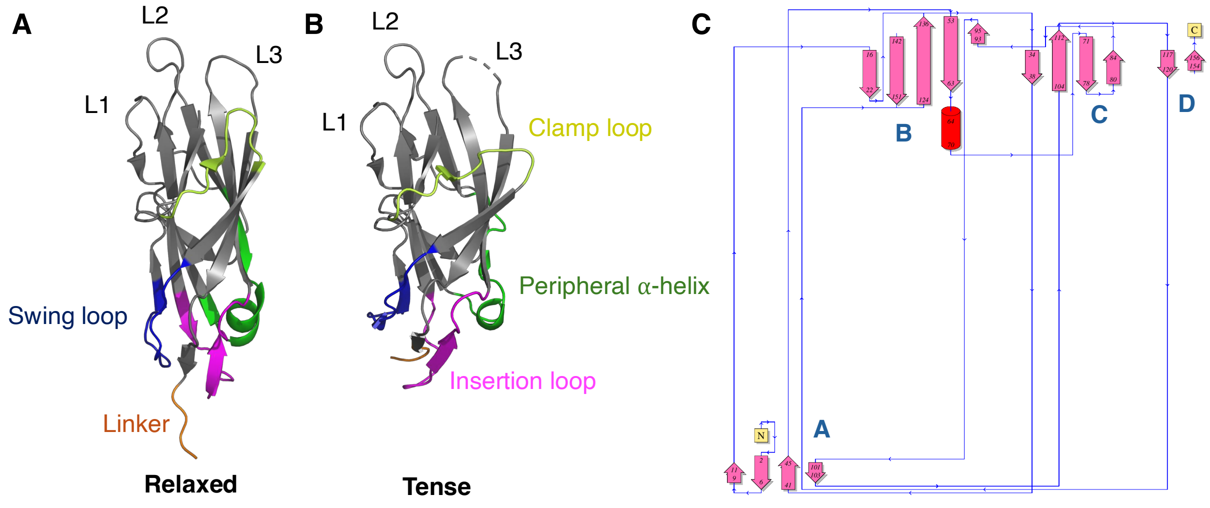

D

```
EcFmlH      FSCNVDGGS--SIGAGTTSVYVNLDPVIQPGQNLVVDLSQHISCWNDYGGWYDTDHINLV       58
EcFimH      FACKTANGTAIPIGGGSANVYVNLAPVVNVGQNLVVDLSTQIFCHNDYPETI-TDYVTLQ       59
KpFimH      FACKTATGATIPIGGGSANVYVNLTPAVNVGQNLVVDLSTQIFCHNDYPETI-TDYVTLQ       59
            *:*:.  *:   **.*::.***** *.:: ********* :* * ***     **::.*

EcFmlH      QGSAFAGSLQSYKGSLYWNNVTYPFPLTTNTNVLDIGDKTPMPLPLKLYITPVGAAGGVV       118
EcFimH      RGSAYGGVLSNFSGTVKYSGSSYPFPTTSETPRVVYNSRTDKPWPVALYLTPVSSAGGVA       119
KpFimH      RGSAYGGVLSSFSGTVKYNGTSYPFPTTTETARVIYDSRTDKPWPAVLYLTPVSTAGGVA       119
            :***:.* *..:.*:: :.. :**** *::*  :  ..:*  * *  **:***.:****.

EcFmlH      IKAGEVIARIHMYKIATLGSGNPRNFTWNIISNNSVVMPTGG  160
EcFimH      IKAGSLIAVLILRQTNNYN-SDDFQFVWNIYANNDVVVPTGG  160
KpFimH      ITAGSLIAVLILHQTNNYN-SDSFQFIWNIYANNDVVVPTGG  160
            *.**.:** : : :  . . .:  :* *** :**.**:****
```

**Fig. S1.**
Comparison of structural regions of $FimH_{LDs}$ as described in (*29*) colored on **(A)** $FimH_{LD}$ in the relaxed conformation (PDB 5jr4) and **(B)** $FimH_{LD}$ in the tense conformation (PDB 5jqi). The regions are colored as follows: clamp loop in limon (residues 8-16), swing loop in blue (residues 22-35), peripheral α-helix in green (residues 59-72), insertion loop in magenta (residues 109-124), and linker to pilin domain in orange (residues 157-160). **(C)** Topology diagram of $FimH_{LD}$ with β-sheet lettered in dark blue. **(D)** Amino acid sequence alignment of E. coli $FmlH_{LD}$ (EcFmlH), E. coli $FimH_{LD}$ (EcFimH), and K. pneumoniae $FimH_{LD}$ (KpFimH).

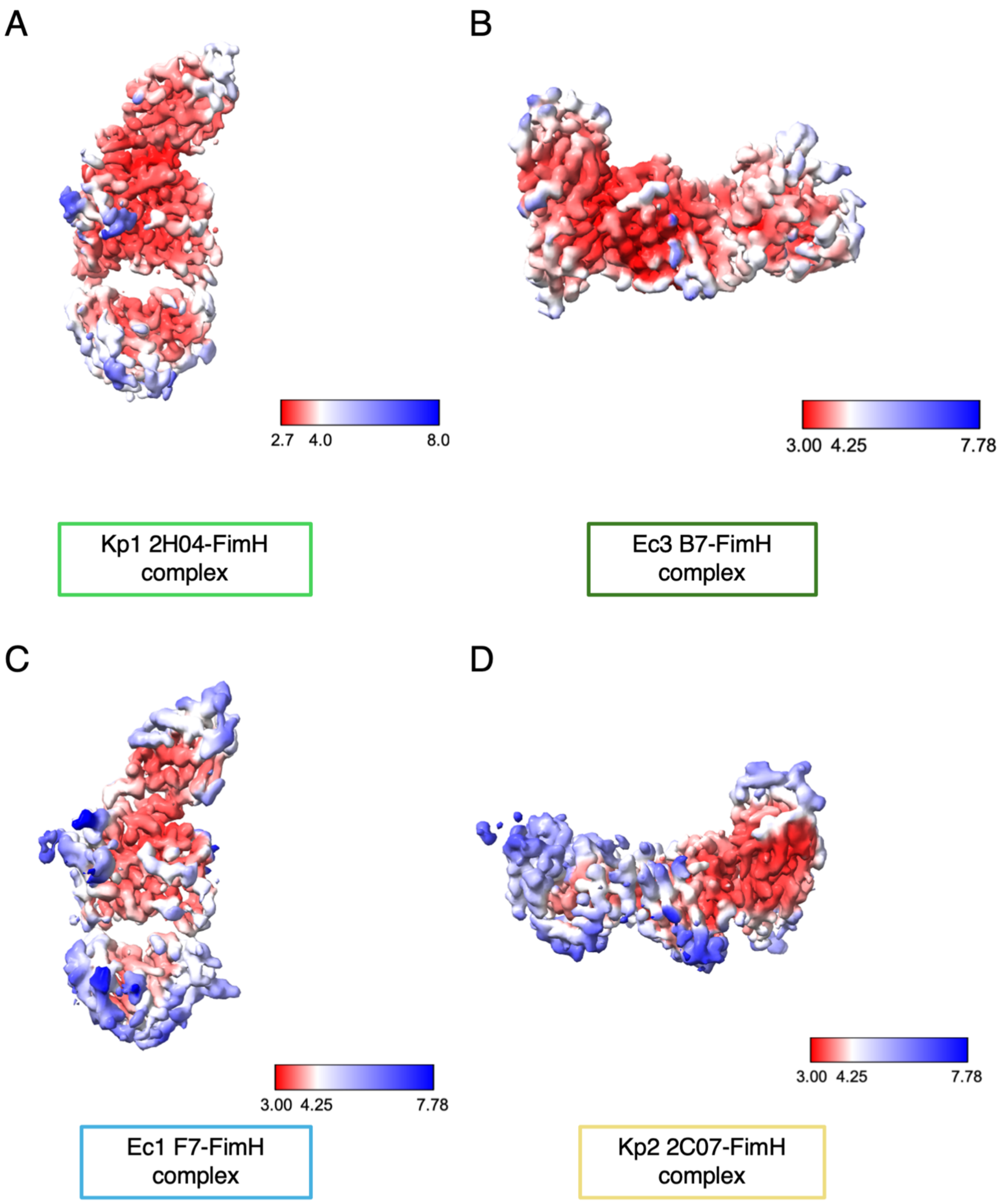

**Fig. S2.**
Local resolution maps of the Fab-FimH structures colored from high resolution (red) to low resolution (blue). **(A)** 2H04-FimH complex, **(B)** B7-FimH complex, **(C)** F7-FimH complex, and **(D)** 2C07-FimH complex. Density maps are viewed at contour level = 0.07.

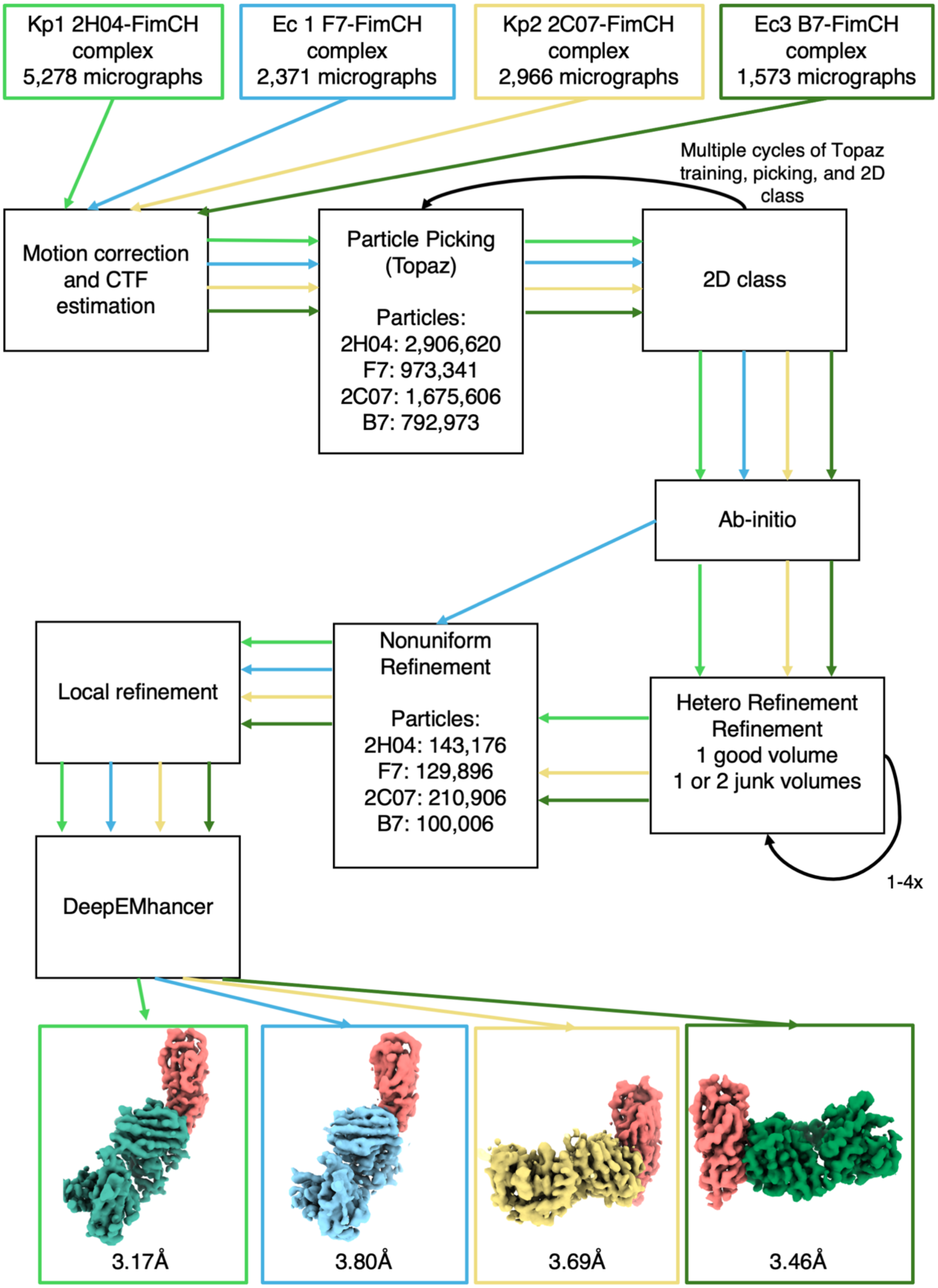

**Fig. S3.**

CryoEM processing flow diagram of Fabs-FimH complexes. For each Fab-FimH complex, data processing is tracked by arrows starting at the top of the diagram and finishing with final density maps at the bottom.

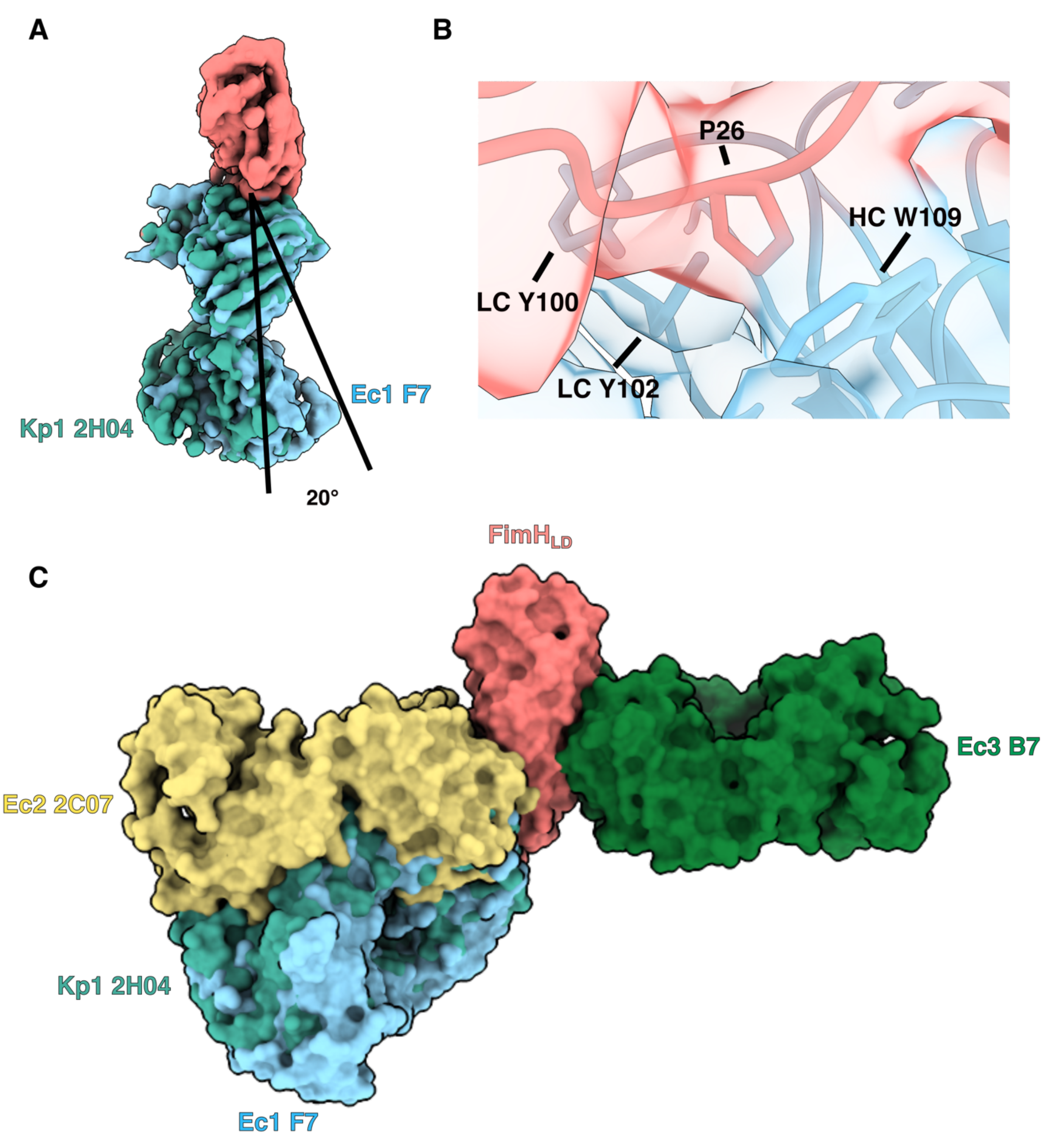

**Fig. S4.**

**(A)** There is a ~20° binding angle shift in the Kp1 2H04 (teal) and Ec1 F7 (cyan) Fabs relative to the binding site on FimH (salmon). Bars represent a 20° angle. **(B)** Ec1 F7 (cyan) coordinated multiple aromatic residues around FimH (salmon). Density map is overlaid on model. mAb heavy chain residues are labeled "HC" and light chain residues are labeled "LC". **(C)** Composite surface models of Kp1 2H04 (teal), Ec1 F7 (cyan), Kp2 2C07 (sand), and Ec3 B7 (green) Fabs on FimH (salmon).

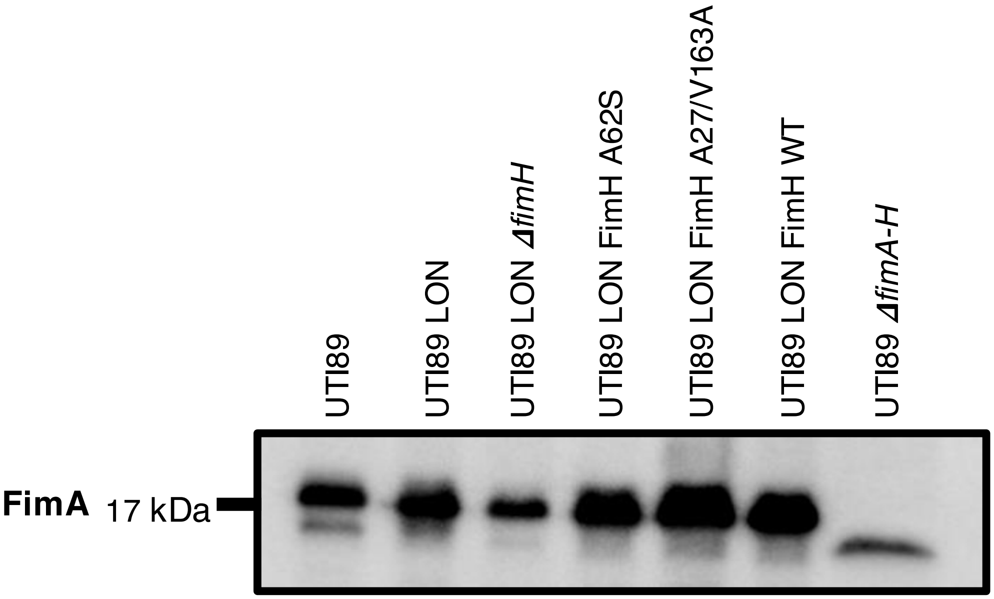

**Fig. S5.**
Western blot of anti-type 1 pili (FimA ~17 kDa) to cell lysates used in bacterial cell ELISA (normalized by $OD_{600}$).

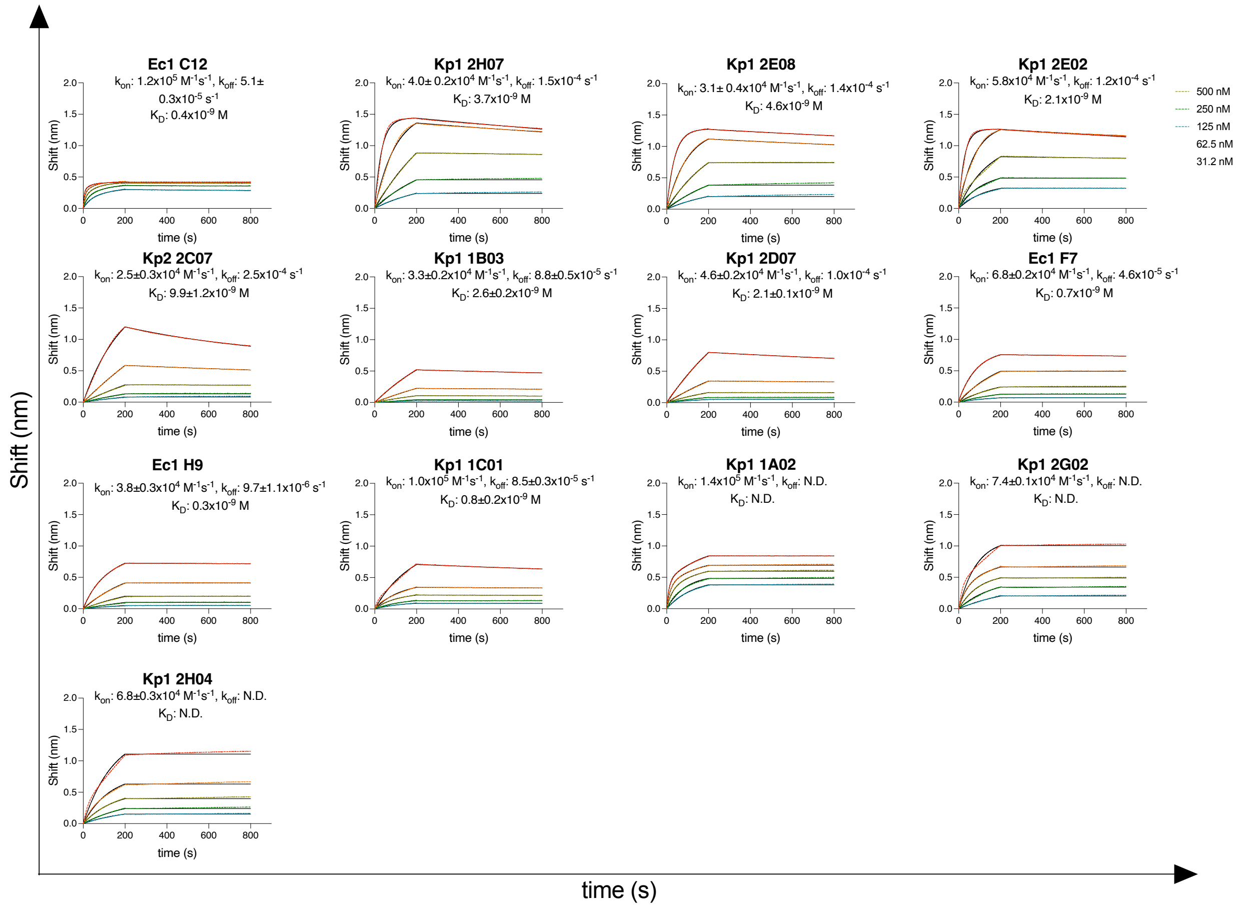

**Fig. S6.**
BLI binding curves of Fabs to *E. coli* $FimG_{nte}H$. Results are from kinetic measurements of dilution series of one experiment.

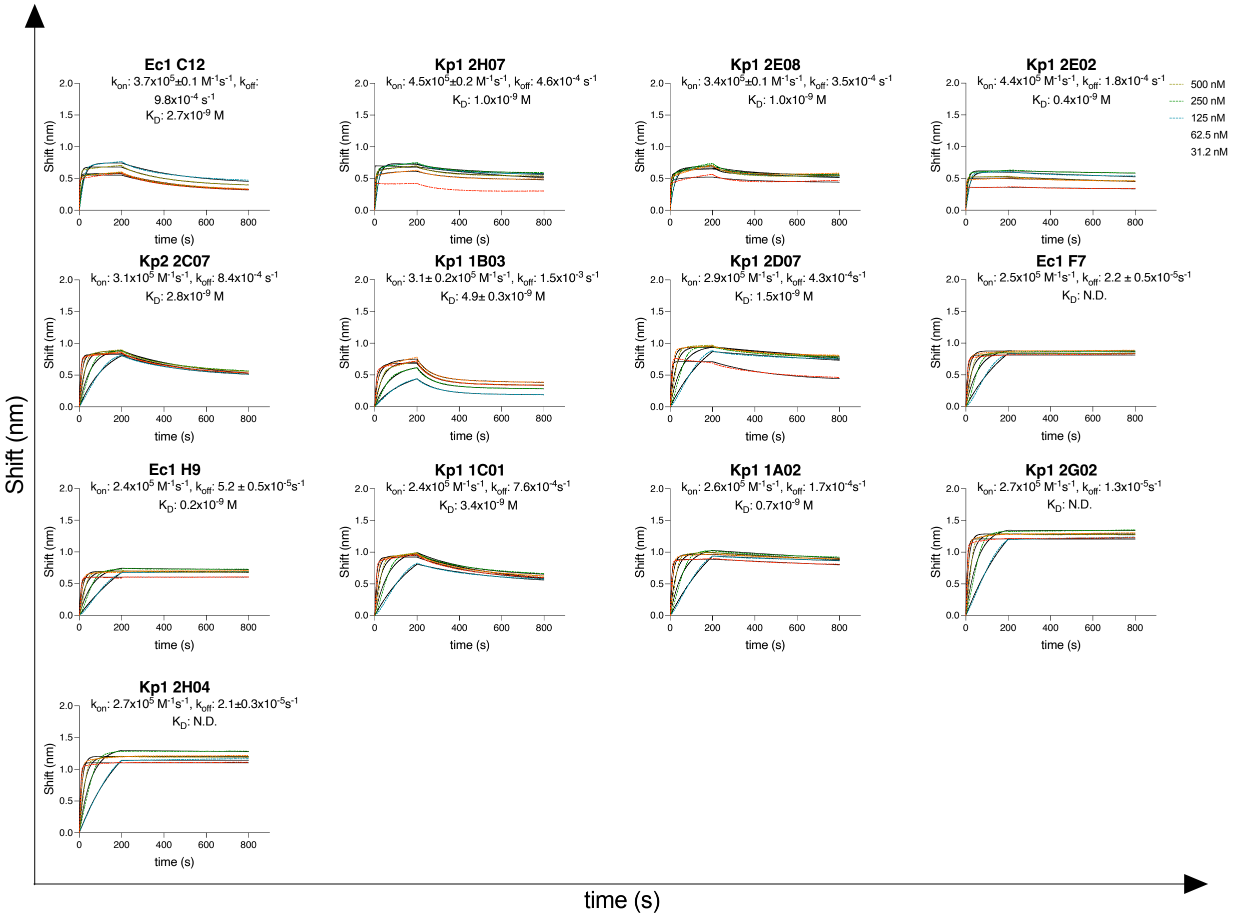

**Fig. S7.**

BLI binding curves to *E. coli* $FimH_{LD}$. Results are from kinetic measurements of dilution series of one experiment.

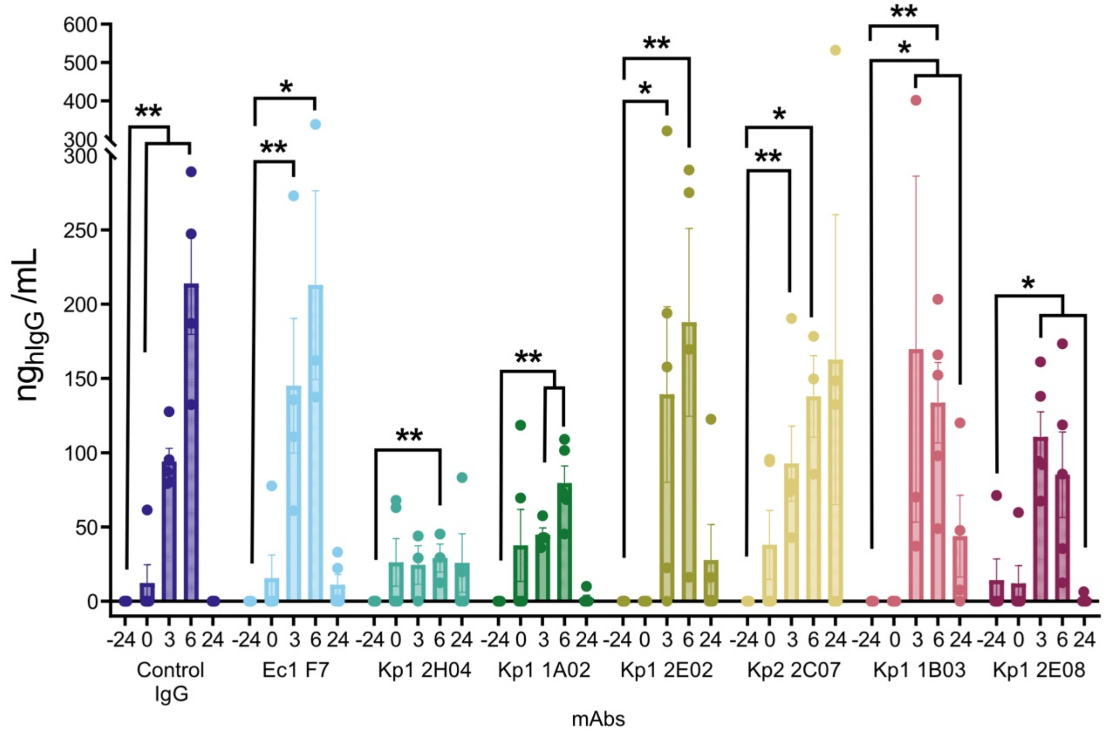

**Fig. S8.**

Presence of FimH mAbs in urine during cystitis. 0.5 mg of selected mAbs were injected via IP. Levels of hIgG mAbs in urine (1:10 dilution) at -24 (Pre-IP), 0 (Pre-infection), 3, 6 and 24 hrs for each of the FimH-specific mAbs and IgG control (n=3 to 5 per group). Mann-Whitney test used to determine significance.

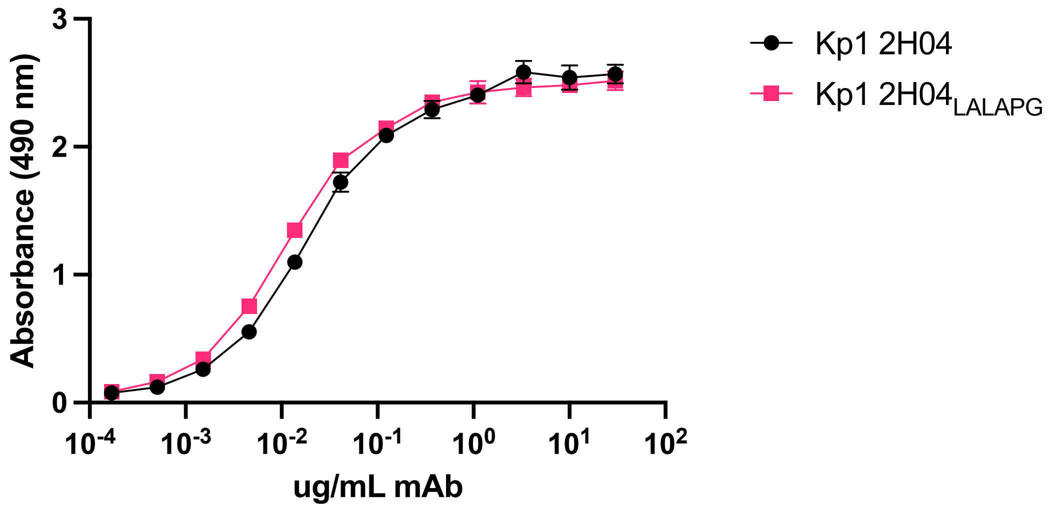

**Fig. S9.**
ELISA binding of Kp1 2H04 and Kp1 $2H04_{LALAPG}$ mAbs to *E. coli* $FimH_{LD}$ (n=3).

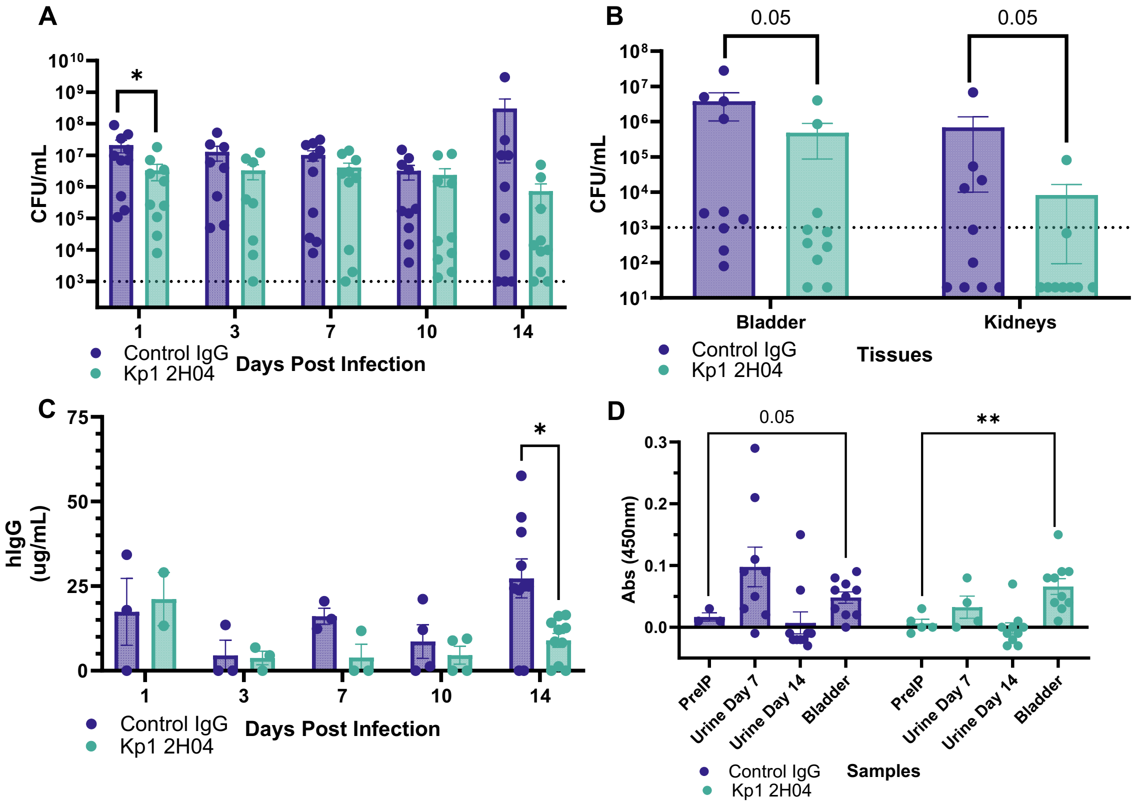

**Fig. S10.**

Protective Kp1 2H04 is detectable in serum two weeks after intraperitoneal injection. Titers in urine **(A)** were measured during the two-week infection period, and bacterial loads in bladder and kidney tissues **(B)** were assessed at sacrifice from mice pretreated with 0.5 mg of Kp1 2H04 (n=10) or control IgG (n=10) 24 hours before infection with UTI89. Horizontal dashed lines represent limit of detection (LOD) of UTI89 titers in urine and tissues. **(C, D)** hIgG mAb levels in serum (**C**, 1:100 dilution), urine (**D**, 1:10 dilution), and bladder (at sacrifice in day 14) homogenates (**D**, 1:2 dilution) of mice pretreated with 0.5 mg of hIgG mAb before infection. Mann-Whitney test was used to determine significance.

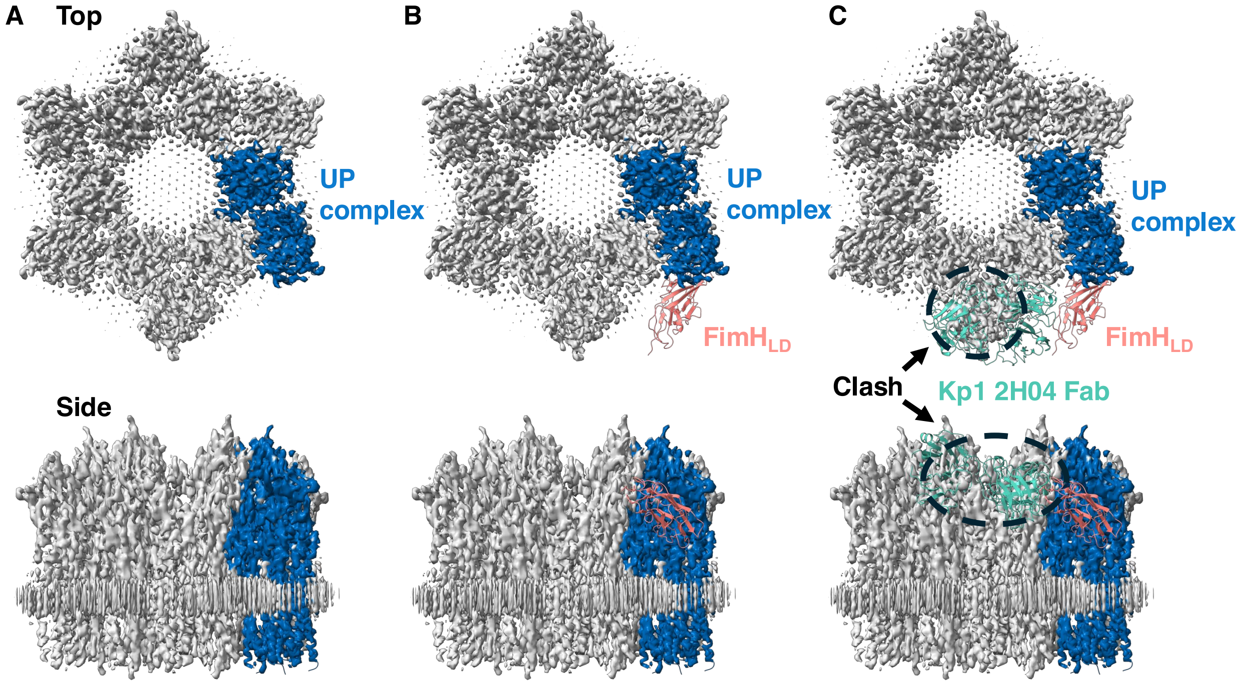

**Fig. S11.**

Top and side view of **(A)** cryoEM density map of singular uroplakin complex (singular unit in blue; EMD-36340) modeled with **(B)** bound FimH lectin domain (salmon) and **(C)** FimH lectin domain-Kp1 2H04 Fab complex (Kp1 2H04 Fab in teal) (*41*, *61*). FimH is modeled to bind to a glycan off N169 as supported by previous evidence (*41*, *61*). Structural clashes between uroplakin complex density and Fab model are noted, despite FimH being in a high-affinity state.

**Table S1.**
mAb class and epitope descriptions between tense and relaxed states.

| Class | mAbs | Lectin domains bound | FimH epitope | FimH regions | Highlighted on Relaxed (blue) structure | Highlighted on Tense (purple) structure |
|---|---|---|---|---|---|---|
| 1 | Ec1 C12<br>Ec1 F11<br>Ec1 F7<br>Ec1 H9<br>Kp1 1A02<br>Kp1 1A09<br>Kp1 1B03<br>Kp1 1C01<br>Kp1 1E07<br>Kp1 2D07<br>Kp1 2D10<br>Kp1 2E02<br>Kp1 2E08<br>Kp1 2G02<br>Kp1 2G04<br>Kp1 2H04<br>Kp1 2H07 | Ec FimH<br>Kp FimH<br>Ec FmlH | 23-29, 32, 118, 121-123, 151-153, 155, 157-159 | Insertion loop, Swing loop, Linker | 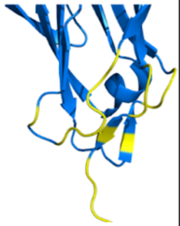 | 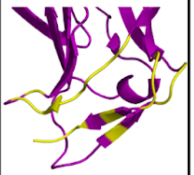 |
| 2 | Kp2 1B02<br>Kp2 2C07<br>Kp2 2C09<br>Kp2 2D04 | Ec FimH<br>Kp FimH | 4, 6, 7, 21, 23-27, 35-37, 40, 41, 43, 122, 123, 151, 152 | Clamp loop, Swing loop | 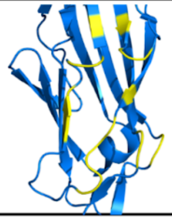 | 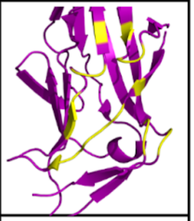 |
| 3 | Ec3 A2<br>Ec3 B7<br>Kp3 2A02<br>Kp3 1D09<br>Kp3 2B06 | Ec FimH<br>Kp FimH | 59, 60, 64, 67-69, 72, 83-89, 121, 125, 128, 130, 132, 143, 145, 147, 155 | β-sheet below clamp loop, Peripheral α-helix | 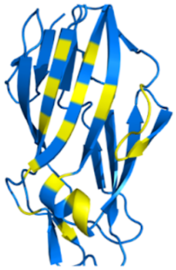 | 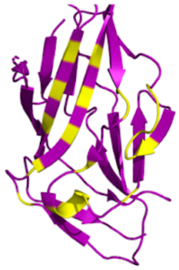 |
| 4 | Ec4 C7<br>Ec4 E7 | Ec FimH | 55, 78, 80, 92, 101 | Base of binding loop 2, Backside | 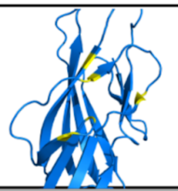 | 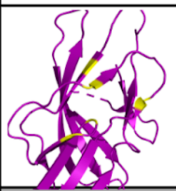 |
| N.D. | Kp 1C08<br>Kp 1B07<br>Kp 1E04<br>Kp 1G01<br>Kp 2A03 | | | | | |

**Table S2.**

Bacterial stains used in this study

| **Strain** | **Reference** |
|---|---|
| C600 ptrc99a-Ec FimH LD | (*45*) |
| C600 ptrc99a-Kp FimH LD | (*21*) |
| C600 ptrc99a-Ec FmlH LD | (*27*) |
| C600 Δ*fim* pBAD33-FimH (J96 WT or mutants for epitope mapping). C600 Δ*fim* was created using standard red recombinase cloning technique. | This study, (*62*) |
| C600 pBAD33-FimH UTI89 WT ptrc99a-FimC | (*42*) |
| C600 pBAD33-FimH UTI89 A62S ptrc99a-FimC | (*42*) |
| UTI89 LON (FimS LIR mutant) | (*63*, *64*) |
| UTI89 | (*14*, *43*) |
| UTI89 LON *ΔfimH* (FimS LIR mutant) | (*63*) |
| UTI89 LON FimH WT (*ΔfimBE*) | (*43*) |
| UTI89 LON FimH A27V/V163A (*ΔfimBE*) | (*43*) |
| UTI89 LON FimH A62S (*ΔfimBE*) | (*43*) |
| UTI89 *ΔfimA-H* | (*15*) |
| UTI89 HK:KanR GFP | (*65*) |
| UTI89 pcomGFP | (*66*) |

**Table S3.**

Primers used for mAb generation

| |
|---|
| **1st Round PCR Primers** |
| IgG, IgK, IgL primers as found in (*26*) |
| IgM/A |
| Forward |
| VH/Outer as found in (*49*): GGGAATTCGAGGTGCAGCTGCAGGAGTCTGG |
| Reverse |
| 3’Cμ outer as found in (*49*): AGGGGGCTCTCGCAGGAGACGAGG |
| 3’Cα outer as found in (*48*): GAAAGTTCACGGTGGTTATATCC |
| **Nested PCR primers** |
| IgG, IgK, IgL primers as found in (*26*) |
| IgM/A |
| Forward |
| VH/Outer as found in (*49*): GGGAATTCGAGGTGCAGCTGCAGGAGTCTGG |
| Reverse |
| 3’Cμ inner as found in (*49*): AGGGGGAAGACATTTGGGAAGGAC |
| 3’Cα inner as found in (*48*): TGCCGAAAGGGAAGTAATCGTGAAT |
| **Gibson cloning primers** |
| IgH |
| Forward |
| VH01: ATCCTTTTTCTAGTAGCAACTGCAACCGGTGTACATTCCGAGGTCCARCTGCARCAGYCTGG |
| VH02: CCTTTTTCTAGTAGCAACTGCAACCGGTGTACATTCCCAGGTGCAGCTGAAGSAGTC |
| VH06: CCTTTTTCTAGTAGCAACTGCAACCGGTGTACATTCCGAAGTGAAGCTTGARGWGTCTG |
| VH14: CCTTTTTCTAGTAGCAACTGCAACCGGTGTACATTCCGAGGTTCAGCTGCAGCAG |
| Reverse |
| JH01: GAAGACCGATGGGCCCTTGGTCGACGCTGAGGAGACGGTGACCGTG |
| JH02: GAAGACCGATGGGCCCTTGGTCGACGCTGAGGAGACTGTGAGA |
| JH03: GAAGACCGATGGGCCCTTGGTCGACGCTGCAGAGACAGTGACCAGAG |
| JH04: GAAGACCGATGGGCCCTTGGTCGACGCTGAGGAGACGGTGACTGAG |
| IgK |

| |
|---|
| Forward |
| VK01: CTTTTTCTAGTAGCAACTGCAACCGGTGTACATTCCGATGTTGTGATGACCCARACTC |
| VK03: CTTTTTCTAGTAGCAACTGCAACCGGTGTACATTCCGACATTGTGCTGACCCAATCTC |
| VK04: CTTTTTCTAGTAGCAACTGCAACCGGTGTACATTCCCAAATTGTTCTCACCCAGTCTC |
| VK05: CTTTTTCTAGTAGCAACTGCAACCGGTGTACATTCCGACATTGTGCTGACYCAGTCTC |
| VK06: CTTTTTCTAGTAGCAACTGCAACCGGTGTACATTCCGACATTGTGATGACCCAGTCTC |
| VK08: CTTTTTCTAGTAGCAACTGCAACCGGTGTACATTCCGACATTGTGATGACMCAGTC |
| VK10 (*67*): CTTTTTCTAGTAGCAACTGCAACCGGTGTACATTCCGACATCCAGATGACTCAGTCTCCA |
| VK12: CTTTTTCTAGTAGCAACTGCAACCGGTGTACATTCCGACATCCAGATGACTCAGTCTC |
| VK14: CTTTTTCTAGTAGCAACTGCAACCGGTGTACATTCCGACATCAAGATGACCCARTCTC |
| Reverse as found in (*67*) |
| 3' Gib-mJK01: AGACAGATGGTGCAGCCACCGTACGTTTGATTTCCAGCTTGGTG |
| 3' Gib-mJK02: AGACAGATGGTGCAGCCACCGTACGTTTTATTTCCAGCTTGGTC |
| 3' Gib-mJK04: AGACAGATGGTGCAGCCACCGTACGTTTTATTTCCAACTTTGTC |
| 3' Gib-mJK05: AGACAGATGGTGCAGCCACCGTACGTTTCAGCTCCAGCTTGGTC |
| IgL (adapted from (*26*)) |
| Forward |
| MLFHLGib_2: ATCCTTTTTCTAGTAGCAACTGCAACCGGTGTACATTCCCAGGCTGTTGTGACTCAG |
| Reverse |
| MLRHLGib_2: TGTTGGCTTGAAGCTCCTCACTCGAGGGYGGAAACAGGGTGACTGATGGCGAAGACTT |
| MLRHLGib_3: TGTTGGCTTGAAGCTCCTCACTCGAGGGYGGAAACACGGTGAGAGTGGGAGTGGACTT |

**Table S4.**
CryoEM model refinement and validation statistics

| | Kp1 2H04-FimH | Ec1 F7-FimH | Ec3 B7-FimH | Kp2 2C07-FimH |
|---|---|---|---|---|
| **Composition (#)** | | | | |
| Chains | 3 | 3 | 3 | 3 |
| Atoms | 4532 (Hydrogens: 0) | 4501 (Hydrogens: 0) | 4491 (Hydrogens: 0) | 4558 (Hydrogens: 0) |
| Residues | Protein: 597 Nucleotide: 0 | Protein: 595 Nucleotide: 0 | Protein: 595 Nucleotide: 0 | Protein: 601 Nucleotide: 0 |
| Water | 0 | 0 | 0 | 0 |
| Ligands | 0 | 0 | 0 | 0 |
| **Bonds (RMSD)** | | | | |
| Length (Å) (# > 4σ) | 0.005 (0) | 0.005 (0) | 0.004 (0) | 0.004 (0) |
| Angles (°) (# > 4σ) | 0.852 (1) | 0.911 (2) | 0.707 (3) | 0.809 (4) |
| MolProbity score | 2.26 | 2.5 | 2.08 | 2.19 |
| Clash score | 15.01 | 15.67 | 11.64 | 12.25 |
| **Ramachandran plot (%)** | | | | |
| Outliers | 0.34 | 0.34 | 0.34 | 0.34 |
| Allowed | 10.83 | 16.81 | 8.15 | 11.43 |
| Favored | 88.83 | 82.85 | 91.51 | 88.24 |
| **Rama-Z (Ramachandran plot Z-score, RMSD)** | | | | |
| whole (N = 591) | -3.25 (0.32) | -3.22 (0.33) | -1.79 (0.35) | -3.03 (0.32) |
| helix (N = 22) | -2.39 (0.70) | -3.52 (0.92) | -1.60 (1.00) | -1.32 (1.92) |

| | | | | |
|---|---|---|---|---|
| sheet (N = 220) | -2.27 (0.30) | -1.22 (0.35) | -0.51 (0.35) | -1.41 (0.36) |
| loop (N = 349) | -2.07 (0.33) | -2.78 (0.32) | -1.72 (0.34) | -2.47 (0.29) |
| Rotamer outliers (%) | 0.78 | 1.38 | 0 | 0 |
| Cβ outliers (%) | 0 | 0 | 0 | 0 |
| Peptide plane (%) | | | | |
| Cis proline/general | 9.1/0.0 | 8.6/0.0 | 8.8/0.0 | 10.3/0.0 |
| Twisted proline/general | 3.0/0.2 | 0.0/0.0 | 0.0/0.0 | 3.4/0.0 |
| CaBLAM outliers (%) | 5.13 | 8.92 | 5.49 | 7.3 |
| **ADP (B-factors)** | | | | |
| Iso/Aniso (#) | 4532/0 | 4501/0 | 4491/0 | 4558/0 |
| min/max/mean | | | | |
| Protein | 18.79/129.80/74.18 | 61.50/146.72/91.65 | 25.70/128.99/62.80 | 49.62/132.53/75.87 |
| Nucleotide | --- | --- | --- | --- |
| Ligand | --- | --- | --- | --- |
| Water | --- | --- | --- | --- |
| **Occupancy** | | | | |
| Mean | 1 | 1 | 1 | 1 |
| occ = 1 (%) | 100 | 100 | 100 | 100 |
| 0 < occ < 1 (%) | 0 | 0 | 0 | 0 |
| occ > 1 (%) | 0 | 0 | 0 | 0 |
| CC (mask) | 0.65 | 0.61 | 0.77 | 0.58 |

| | | | | |
|---|---|---|---|---|
| CC (box) | 0.67 | 0.66 | 0.78 | 0.66 |
| CC (peaks) | 0.64 | 0.61 | 0.77 | 0.59 |
| CC (volume) | 0.65 | 0.61 | 0.78 | 0.58 |
| Resolution (Å) | 3.2 | 3.8 | 3.5 | 3.7 |

**Table S5.**
Primers used for FimH mutatgenesis

| AA | Codon | Mutant Codon | Primer Name | Primer Sequence |
|---|---|---|---|---|
| pBAD | N/A | N/A | pBAD33 F | ATGCCATAGCATTTTTATCC |
| | | | pBAD33 R | GATTTAATCTGTATCAGG |
| 4K | AAA (K) | GAT (D) | J96_FimH 4K F | GCCTGTGATACCGCCAATGGTACC |
| | | | J96_FimH 4K R | GGCGGTATCACAGGCGAATGACC |
| 7N | AAT (N) | AAA (K) | J96_FimH 7N F | GCCTGTAAAACCGCCAAAGGTACC |
| | | | J96_FimH 7N R | GATAGCGGTACCTTTGGCGGTTTT |
| 10A | GCT (A) | GAT (D) | J96_FimH 10A F | GGTACCGATATCCCTATTGGCGGTG |
| | | | J96_FimH 10A R | GCCAATAGGGATATCGGTACCATTGGC |
| 13I | ATT (I) | AAA (K) | J96_FimH 13I F | GCTATCCCTAAAGGCGGTGGCAGC |
| | | | J96_FimH 13I R | CCACCGCCTTTAGGGATAGCGG |
| 17S | AGC (S) | AAA (K) | J96_FimH 17S F | GGTGGCAAAGCCAATGTTTATGTAAACCTTGCG |

| | | | | |
|---|---|---|---|---|
| | | | J96_FimH 17S R | GTTTACATAAACATTGGCTTTGCCACCGCCAATAGG |
| 19N | AAT (N) | AAA (K) | J96_FimH 19N F | GCAGCGCCAAAGTTTATGTAAACCTTGC |
| | | | J96_FimH 19N R | GTTTACATAAACTTTGGCGCTGCCACC |
| 21Y | TAT (Y) | GAT (D) | J96_FimH 21Y F | GCCAATGTTGATGTAAACCTTGCGCCC |
| | | | J96_FimH 21Y R | GTTTACATCAACATTGGCGCTGCCACC |
| 23N | AAC (N) | AAA (K) | J96_FimH 23N F | GTTTATGTAAAACTTGCGCCCGTCGTG |
| | | | J96_FimH 23N R | GACGGGCGCAAGTTTTACATAAACATTGGC |
| 25A | GCG (A) | GAT (D) | J96_FimH 25A-2 F | GTAAACCTTGATCCCGTCGTGAATGTGGGG |
| | | | J96_FimH 25A-2 R | CACGACGGGATCAAGGTTTACATAAACATTGGC |
| 27V | GTC (V) | GAC (D) | J96_FimH 27V F | CCTTGCGCCCGACGTGAATG |
| | | | J96_FimH 27V R | CCACATTCACGTCGGGCGC |
| 29N | AAT (N) | AAA (K) | J96_FimH 29N F | GCCCGTCGTGAAAGTGGGG |
| | | | J96_FimH 29N R | GGTTTTGCCCCACTTTCACGAC |
| 30V | GTG (V) | GAT (D) | J96_FimH 30V F | CCCGTCGTGAATGATGGGCAAAAC |
| | | | J96_FimH 30V R | CGACCAGGTTTTGCCCATCATTCAC |
| 33N | AAC (N) | AAA (K) | J96_FimH 33N F | CGTGAATGTGGGGCAAAAACTGGTCG |
| | | | J96_FimH 33N R | CGAAAGATCCACGACCAGTTTTTGCCC |

| | | | | |
|---|---|---|---|---|
| 37D | GAT (D) | AAA (K) | J96_FimH 37D F | CCTGGTCGTGAAACTTTCG |
| | | | J96_FimH 37D R | GCGTCGAAAGTTTCACGAC |
| 40T | ACG (T) | AAG (K) | J96_FimH 40T F | CGTGGATCTTTCGAAGCAAATC |
| | | | J96_FimH 40T R | GGCAAAAGATTTGCTTCGAAAG |
| 43F | TTT (F) | GAT (D) | J96_FimH 43F F | CGCAAATCGATTGCCATAACGATT |
| | | | J96_FimH 43F R | GGCAATCGATTTGCGTCGAAAGAT |
| 48Y | TAT (Y) | GAT (D) | J96_FimH 48Y F | GCCATAACGATGATCCGGAAACC |
| | | | J96_FimH 48Y R | CTGTAATGGTTTCCGGATCATCGTTATGG |
| 50E | GAA ( E) | AAA (K) | J96_FimH 50E F | GCCATAACGATTATCCGAAAACCATTAC |
| | | | J96_FimH 50E R | GTCTGTAATGGTTTTCGGATAATCG |
| 55Y | TAT (Y) | GAT (D) | J96_FimH 55Y F | CCATTACAGACGATGTCACACTGC |
| | | | J96_FimH 55Y R | GTGACATCGTCTGTAATGGTTTCC |
| 60R | CGA ( R) | AAA (K) | J96_FimH 60R F | CACTGCAAAAAGGCTCGGCTTATGGCGG |
| | | | J96_FimH 60R R | GCCGAGCCTTTTTGCAGTGTGACATAGTC |
| 62S | TCG (S) | AAG (K) | J96_FimH 62S F | GCAACGAGGCAAGGCTTATGGC |
| | | | J96_FimH 62S R | GCCGTTCCTCGTTGCAGTGTGACA |
| 64Y | TAT (Y) | GAT (D) | J96_FimH 64Y F | CTCGGCTGATGGCGGCGTGTTAT |

| | | | | |
|---|---|---|---|---|
| | | | J96_FimH 64Y R | CCATCAGCCGAGCCTCGTTGCA |
| 67V | GTG (V) | GAT (D) | J96_FimH 67V F | GGCGGCGATTTATCTAATTTTCCGGG |
| | | | J96_FimH 67V R | TTAGATAAATCGCCGCCATAAGCCGAG |
| 70N | AAT (N) | AAA (K) | J96_FimH 70N F | GTTATCTAAATTTTCCGGGACCG |
| | | | J96_FimH 70N R | GGAAAATTTAGATAACACGCCGC |
| 74T | ACC (T) | AAA (K) | J96_FimH 74T F | CCGGGAAAGTAAAATATAGTGGC |
| | | | J96_FimH 74T R | CTATATTTTACTTTCCCGGAAAAATTAG |
| 76K | AAA (K) | GAT (D) | J96_FimH 76K F | CCGGGACCGTAGATTATAGTGGCAGTAGC |
| | | | J96_FimH 76K R | GCCACTATAATCTACGGTCCCGGAAAAAT TAG |
| 78N | AGT (S) | AAA (K) | J96_FimH 78N F | CGTAAAATATAAAGGCAGTAGCTATCC |
| | | | J96_FimH 78N R | CTGCCTTTATATTTTACGGTCCC |
| 80S | AGT (S) | AAA (K) | J96_FimH 80S F | GTGGCAAAAGCTATCCATTTCC |
| | | | J96_FimH 80S R | GGATAGCTTTTGCCACTATATTTTACGG |
| 87T | ACC (T) | AAA (K) | J96_FimH 87T F | CCTACCAAAAGCGAAACGCCG |
| | | | J96_FimH 87T R | CGTTTCGCTTTTGGTAGGAAATGG |
| 89E | GAA ( E) | AAA (K) | J96_FimH 89E F | CCTACCAAAAGCGAAACGCCGC |
| | | | J96_FimH 89E R | CGGCGTTTTGCTGGTGGTAGGA |

| | | | | |
|---|---|---|---|---|
| 92R | CGC ( C) | GAC (D) | J96_FimH 92R F | CGCCGGACGTTGTTTATAATTCG |
| | | | J96_FimH 92R R | TAAACAACGTCCGGCGTTTCGC |
| 96N | AAT (N) | AAA (K) | J96_FimH 96N F | GCGTTGTTTATAAATCGAGAACGG |
| | | | J96_FimH 96N R | CCGTTCTCGATTTATAAACAACGCG |
| 98R | AGA ( R) | AAA (K) | J96_FimH 98R F | GTTTATAATTCGAAAACGGATAAGCCG |
| | | | J96_FimH 98R R | GCTTATCCGTTTTCGAATTATAAACAACG |
| 99T | ACG (T) | AAG (K) | J96_FimH 99T F | GTTTATAATTCGAGAAAGGATAAGCCG |
| | | | J96_FimH 99T R | CGGCTTATCCTTTCTCGAATTATAAAC |
| 101K | AAG (K) | GAT (D) | J96_FimH 101K F | GAGAACGGATGATCCGTGGCCGGTG |
| | | | J96_FimH 101K R | CCGGCCACGGATCATCCGTTCTCGAA |
| 110T | ACG (T) | AAG (K) | J96_FimH 110T F | GGTGGCGCTTTATTTGAAGCCTGTGAGC |
| | | | J96_FimH 110T R | CGCACTGCTCACAGGCTTCAAATAAAGC |
| 121K | AAA (K) | GAT (D) | J96_FimH 121K F | GGCGATTGATGCTGGCTCATTAATTGC |
| | | | J96_FimH 121K R | GAGCCAGCATCAATCGCCACC |
| 128V | GTG (V) | GAT (D) | J96_FimH 128V F | CATTAATTGCCGATCTTATTTTGCGAC |
| | | | J96_FimH 128V R | CAAAATAAGATCGGCAATTAATGAGCC |
| 132R | CGA ( R) | AAA (K) | J96_FimH 132R F | GCTTATTTTGAAACAGACCAACAAC |

| | | | | |
|---|---|---|---|---|
| | | | J96_FimH 132R R | GGTCTGTTTCAAAATAAGCACGGC |
| 139S | AGC (S) | AAA (K) | J96_FimH 139S F | CAACTATAACAAAGATGATTTCCAG |
| | | | J96_FimH 139S R | GGAAATCATCTTTGTTATAGTTGTTGG |
| 142F | TTC (F) | GAC (D) | J96_FimH 142F F | CTATAACAGCGATGATGACCAGTTTGTGTGG |
| | | | J96_FimH 142F R | CACAAACTGGTCATCATCGCTGTTATAGTTG |
| 145V | GTG (V) | GAT (D) | J96_FimH 145V F | TTTCCAGTTTGATTGGAATATTTACG |
| | | | J96_FimH 145V R | GTAAATATTCCAATCAAACTGGAAATC |
| 152N | AAT (N) | AAA (K) | J96_FimH 152N F | CGCCAATAAAGATGTGGTGGTG |
| | | | J96_FimH 152N R | CACCACATCTTTATTGGCGTAA |
| 155V | GTG (V) | GAT (D) | J96_FimH 155V F | CCAATAATGATGTGGATGTGCCTACTGGC |
| | | | J96_FimH 155V R | GTAGGCACATCCACATCATTATTGGCG |
